# Cryo-EM structure of the SARS-CoV-2 3a ion channel in lipid nanodiscs

**DOI:** 10.1101/2020.06.17.156554

**Authors:** David M. Kern, Ben Sorum, Sonali S. Mali, Christopher M. Hoel, Savitha Sridharan, Jonathan P. Remis, Daniel B. Toso, Abhay Kotecha, Diana M. Bautista, Stephen G. Brohawn

## Abstract

Severe acute respiratory syndrome coronavirus 2 (SARS-CoV-2) is the virus that causes the coronavirus disease 2019 (COVID-19). SARS-CoV-2 encodes three putative ion channels: E, 8a, and 3a^1,2^. 3a is expressed in SARS patient tissue and anti-3a antibodies are observed in patient plasma^3–6^. 3a has been implicated in viral release^7^, inhibition of autophagy^8^, inflammasome activation^9^, and cell death^10,11^ and its deletion reduces viral titer and morbidity in mice^1^, raising the possibility that 3a could be an effective vaccine or therapeutic target^3,12^. Here, we present the first cryo-EM structures of SARS-CoV-2 3a to 2.1 Å resolution and demonstrate 3a forms an ion channel in reconstituted liposomes. The structures in lipid nanodiscs reveal 3a dimers and tetramers adopt a novel fold with a large polar cavity that spans halfway across the membrane and is accessible to the cytosol and the surrounding bilayer through separate water- and lipid-filled openings. Electrophysiology and fluorescent ion imaging experiments show 3a forms Ca^2+^-permeable non-selective cation channels. We identify point mutations that alter ion permeability and discover polycationic inhibitors of 3a channel activity. We find 3a-like proteins in multiple *Alphacoronavirus* and *Betacoronavirus* lineages that infect bats and humans. These data show 3a forms a functional ion channel that may promote COVID-19 pathogenesis and suggest targeting 3a could broadly treat coronavirus diseases.

## Main Text

Coronavirus disease 2019 (COVID-19), caused by the SARS-CoV-2 virus, is an ongoing global pandemic. Vaccine and therapeutic development are predominantly focused on the essential virus-encoded Spike, main protease, and RNA-dependent RNA polymerase proteins. High-resolution structures of these targets, some in complex with drug candidates or neutralizing antibodies, have yielded mechanistic insight into their function and have provided a platform for successful structure-guided vaccine and drug design^13–18^. Expanding our knowledge of SARS-CoV-2 target proteins remains important both for understanding SARS-CoV-2 virology and developing alternative treatments to mitigate against potential resistance that evolves, or new viruses that emerge, in the future^19^.

The SARS-CoV-2 genome encodes three putative ion channels (viroporins)^20,21^, the envelope protein (E) and open reading frame (ORF) 3a and 8a^1,2^. Viroporins have been shown to modify host membrane permeability to promote viral assembly and release^20,22^ and SARS-CoV-1 3a has been reported to form an emodin-sensitive K^+^-permeable cation channel^23,24^. 3a is highly conserved within the *Betacoronavirus* subgenus *Sarbecovirus* which includes SARS-CoV-1 and other related bat coronaviruses (Fig. S1)^25^. Biopsies from SARS-CoV-1 patients show 3a expression in infected tissue and SARS-CoV-1 and SARS-CoV-2 patient plasma contain anti-3a antibodies^3–6^. SARS-CoV-1 3a has been implicated in inflammasome activation^9^ and both apoptotic^10^ and necrotic cell death^11^ while SARS-CoV-2 3a has been implicated in apoptosis and inhibition of autophagy *in vitro*^26^. In mouse models of SARS-CoV-1 infection, genomic deletion of 3a reduced viral titer and morbidity^1^. 3a has therefore been considered a potential target for vaccines or therapeutics to treat SARS^3,12^.

Still, the precise roles of 3a in disease pathogenesis are unknown, precluded in part by the lack of a mechanistic understanding of 3a. Here, we utilize cryo-electron microscopy (cryo-EM), electrophysiology, and fluorescent ion flux assays to elucidate 3a protein structure and function.

### Structure determination

We determined structures of dimeric and tetrameric SARS-CoV-2 3a in lipid nanodiscs. Full length SARS-CoV-2 3a was heterologously expressed in *Spodoptera frugiperda* (Sf9) cells with a cleavable C-terminal GFP tag. Purification of 3a in detergent resulted in two species separable by gel filtration (Fig. S2). A majority of 3a runs at a position consistent with a 62 kDa dimer (Fig. S2A,C,E) and ~10-30% runs as a 124 kDa tetramer (Fig. S2). A similar degree of tetramer formation was observed at low concentrations of 3a by fluorescence size-exclusion chromatography, indicative of a biochemically stable species rather than concentration-dependent nonspecific aggregation (Fig. S2E). These data are consistent with previous reports of dimeric and tetrameric SARS-CoV-1 3a observed by western blot^11,23^.

We separately reconstituted dimeric and tetrameric SARS-CoV-2 3a into nanodiscs made from the scaffold protein MSP1E3D1 and a mixture of DOPE, POPC, and POPS lipids and determined their structures by cryo-EM to 2.9 Å and 6.5 Å resolution (Fig. 1,2, S2-6). We also determined the structure of dimeric 3a in the presence of 100uM emodin to 3.7 Å, but observed no significant structural changes from dimeric apo 3a or any indication of bound emodin (Fig. S7-9). Based on a recent report of improved resolution for structures of apoferritin and GABA_A_ using new cryo-EM instrumentation^27^, we asked whether similar improvements would be possible for the sub-100 kDa membrane protein 3a. Indeed, a reconstruction generated from data collected on this new instrumentation, using the same batch of grids, was significantly improved at 2.1 Å (Fig. S10-12).

**Figure 1.**
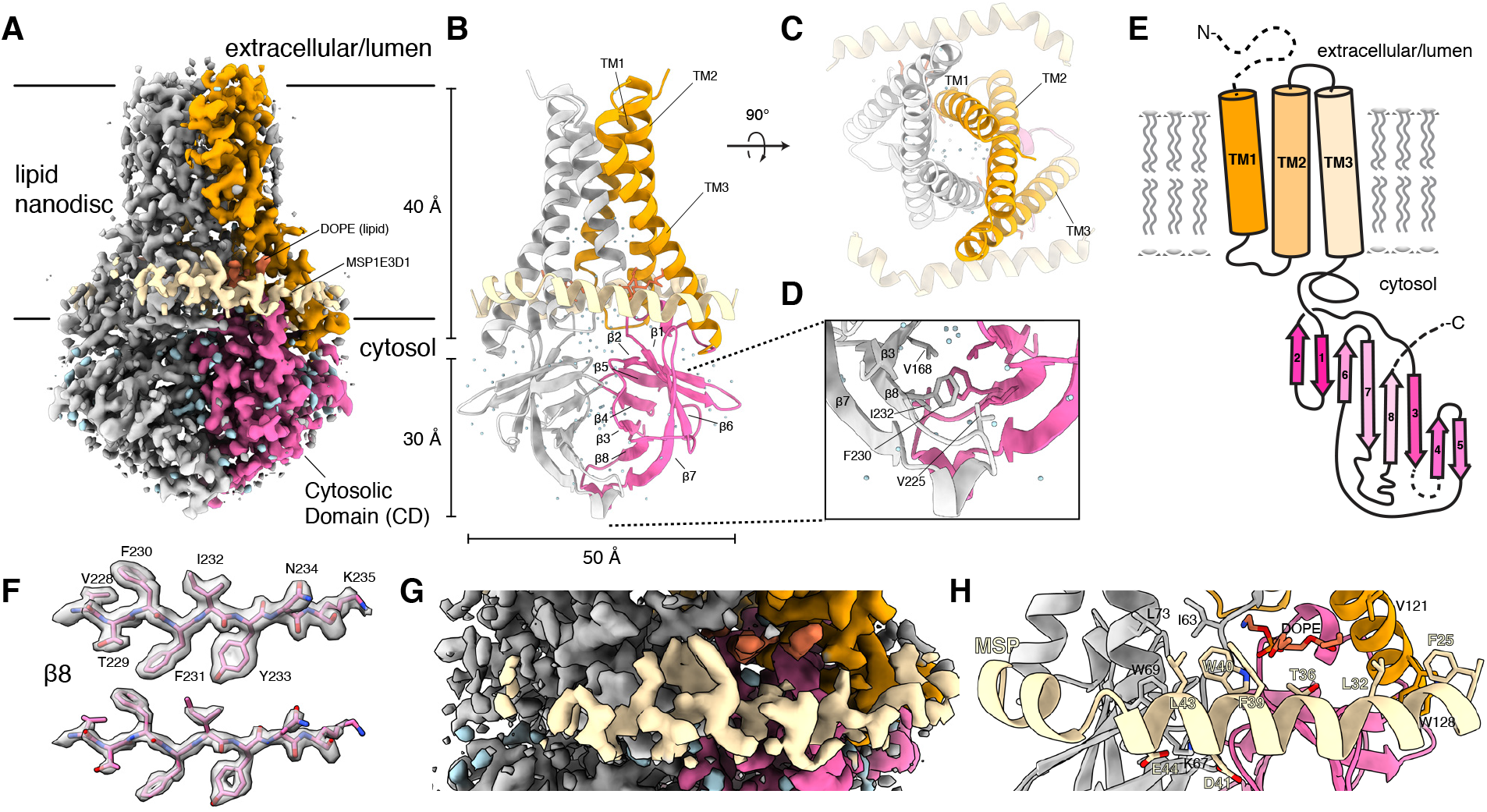
Structure of dimeric 3a in lipid nanodiscs. (A) Cryo-EM map of the 3a dimer in MSP1E3D1 nanodiscs at 2.1 Å nominal resolution viewed from the membrane plane. One subunit is colored gray and the second subunit is colored with transmembrane region orange and cytosolic domain (CD) pink. Density from the nanodisc MSP1E3D1 is colored tan and the DOPE lipid is coral. (B,C) Model of dimeric 3a viewed (B) from the membrane (as in (A)) and (C) from the extracellular or lumenal side. (D) Zoomed in view of the interaction between subunits in the CD with residues forming the hydrophobic core indicated. (E) Schematic of a 3a monomer. Secondary structure elements are indicated and unmodeled termini and a 5 amino acid β3-β4 loop are shown with dashed lines. (F) Cryo-EM density and model for a selected strand from the CD at two different thresholds. (G) Zoomed-in view of density in the MSP1E3D1 interaction region. (H) Model in the same region as H with key residues in the area displayed as sticks.

**Figure 2.**
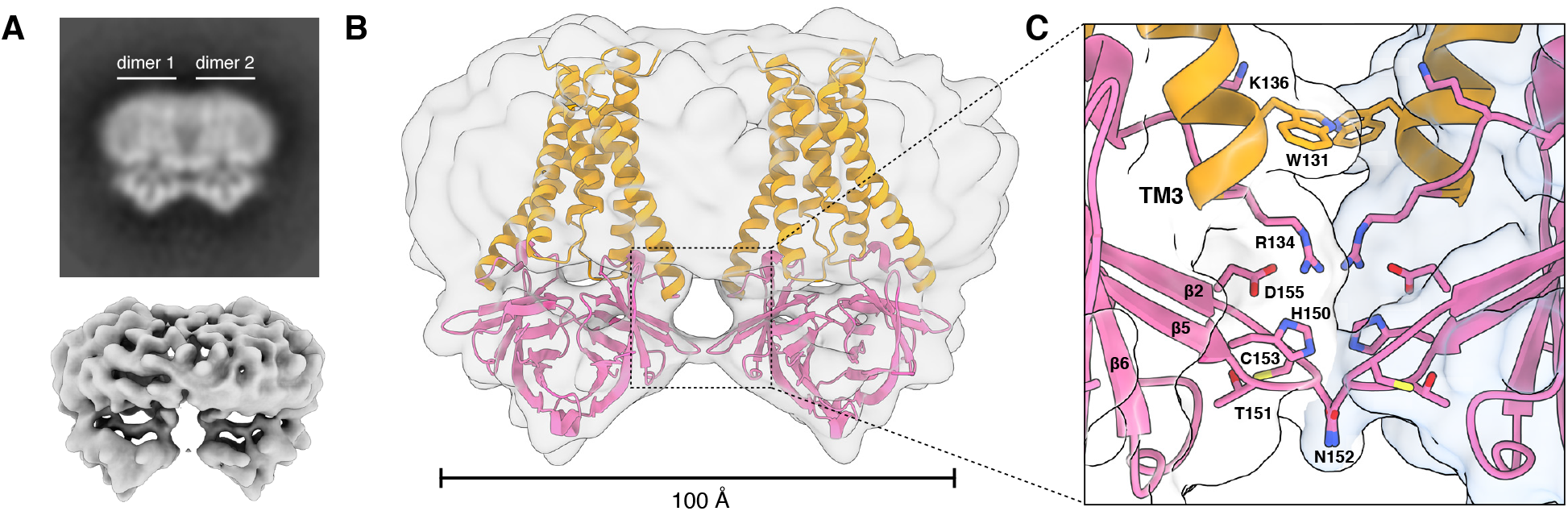
Structure of tetrameric 3a in lipid nanodiscs. (A) Two-dimensional class average of tetrameric 3a in MSP1E3D1 lipid nanodiscs (above) and cryo-EM map at 6.5 Å nominal resolution (lower). (B) Two copies of the dimeric 3a structure rigid-body docked into the tetrameric 3a cryo-EM map. (C) Zoomed in view of the interface between dimers with key residues in the area displayed as sticks.

### Unique features of 3a dimers

The 2.9 Å dimeric reconstruction (with C2 symmetry applied, PDB 6XDC) permitted de novo modeling of 195 of the 275 amino acids per protomer chain. The N-terminus (amino acids 1-39), C-terminus (amino acids 239-275), and a short cytoplasmic loop (amino acids 175-180) are either not observed or weakly resolved in the density map due to conformational differences between particles or because they are disordered (Fig. 1A-E). The 2.1 Å dimeric reconstruction (with C2 symmetry applied, PDB 7KJR) allowed for improved rotamer assignments due to better defined side chain density throughout the protein, including visible holes in many aromatic residues (Fig. 1F). We additionally were able to model a portion of each MSP1E3D1 scaffold protein (amino acids 25-55, Fig. 1G,H), two DOPE lipids, and 122 water molecules. A portion of the scaffold protein is well-resolved largely due to a specific interaction visible on each side of the 3a dimer (even in C1 reconstructions) between MSP1E3D1 W40 and the transmembrane region of 3a (Fig. 1F,G). The symmetric nature of this interaction means that the two MSPE3D1 protomers must twist around the lipid bilayer rather than adopting the canonical arrangement of two parallel stacked rings. We refer to the higher resolution structure unless otherwise indicated.

Surprisingly, 3a adopts a novel fold that, to our knowledge, has not been observed in available protein structures. Querying the protein structure database for homologs with Dali returned only weak hits for fragments of 3a domains^28^. Viewed from the membrane plane, 3a is approximately 70 Å tall with a 40 Å high transmembrane region and a cytosolic domain (CD) extending 30 Å out of the membrane (Fig. 1A,B). The transmembrane region is composed of three helices per protomer. 3a is a Class IIIA viroporin^22^, with N-termini oriented on the extracellular or lumenal side and C-termini on the cytosolic side of the membrane. Viewed from the extracellular side, the transmembrane helices (TMs) trace the circumference of an ellipse with TM1-3 from one protomer followed by TM1-3 of the second protomer in a clockwise order (Fig. 1C). TM1s and TM2s pack against each other across the elliptical minor axis with TM3s positioned at the major axis vertices. TM1-TM2 and TM2-TM3 are joined by short intracellular and extracellular linkers, respectively.

The transmembrane region connects to the CD through a turn-helix-turn motif following TM3. Each protomer chain forms a pair of opposing β-sheets packed against one another in an eight stranded β-sandwich (Fig. 1B,D). The outer sheet is formed by strands β1, β2, β6 and the N-terminal half of β7. The inner sheet is formed by strands β3, β4, β5, β8, and C-terminal half of β7. The inner sheets from each protomer interact through a large (~940 Å^2^ buried surface area per chain) and highly complementary interface with residues V168, V225, F230, and I232 forming a continuous hydrophobic core (Fig. 1D). The interaction between β-sandwiches from each protomer thus forms a strong and stable link between monomers in the dimer.

### Structural features of 3a tetramers

We next examined the structure of 3a tetramers. Two-dimensional class averages of tetrameric 3a show a side by side arrangement of two dimers with well-separated TMs and close juxtaposition of CDs (Fig. 2A). Tetrameric 3a reconstructions had lower final resolutions (~6.5 Å) than dimeric 3a (Fig. 2A, S6), but were sufficiently featured in the CDs to enable rigid-body docking of two copies of the 3a dimer (Fig. 2B). The best fit models show that TM3-CTD linkers and β1-β2 linkers from neighboring dimers form a continuous interface (~300 Å^2^ buried surface area per dimer). While the exact positions of side chains cannot be determined at this resolution, residues W131, R134, K136, H150, T151, N152, C153, and D155 are poised to mediate tetramerization through a network of hydrophobic, polar, and electrostatic interactions (Fig. 2C). In SARS-CoV-1 3a, reducing agents and a C133A mutation resulted in loss of oligomerization, membrane localization, and channel activity^23^. However, expression of the C133A mutant was dramatically reduced, suggesting these results may be a consequence of protein destabilization^23^. In SARS-CoV-2 3a, C133 is located in a notable cysteine-rich pocket adjacent to the tetramerization interface (Fig. S13). At the base of TM3, C133 projects back towards the top faces of β1 and β2 in close proximity to solvent exposed C148 and buried C157. All three cysteines are reduced in the structure, though they are within potential disulfide-bonding distance (Cα distance 4.4-6.5 Å) (Fig. S13). While it is unlikely a disulfide involving C133 mediates tetramerization of 3a without significant rearrangement of this region, it may be that disruption of this cysteine-rich pocket with cysteine modifying agents or mutations disfavors 3a oligomerization allosterically.

### Ion permeation through 3a

Ion conduction pathways are key determinants of ion channel function. Examination of the 3a structure for possible conduction pathways shows a hydrophobic seal in the extracellular half of the transmembrane region above a large polar cavity (Fig. 3). The hydrophobic seal is formed by interactions between transmembrane helices and includes residues F43, L46, I47, V50, L95, and L96 (Fig. 3A-C). Just cytoplasmic to the hydrophobic constriction, the top of the channel cavity is formed by polar interactions between Q57, S58, N82, and Q116 side chains (Fig. 3A,D).

**Figure 3.**
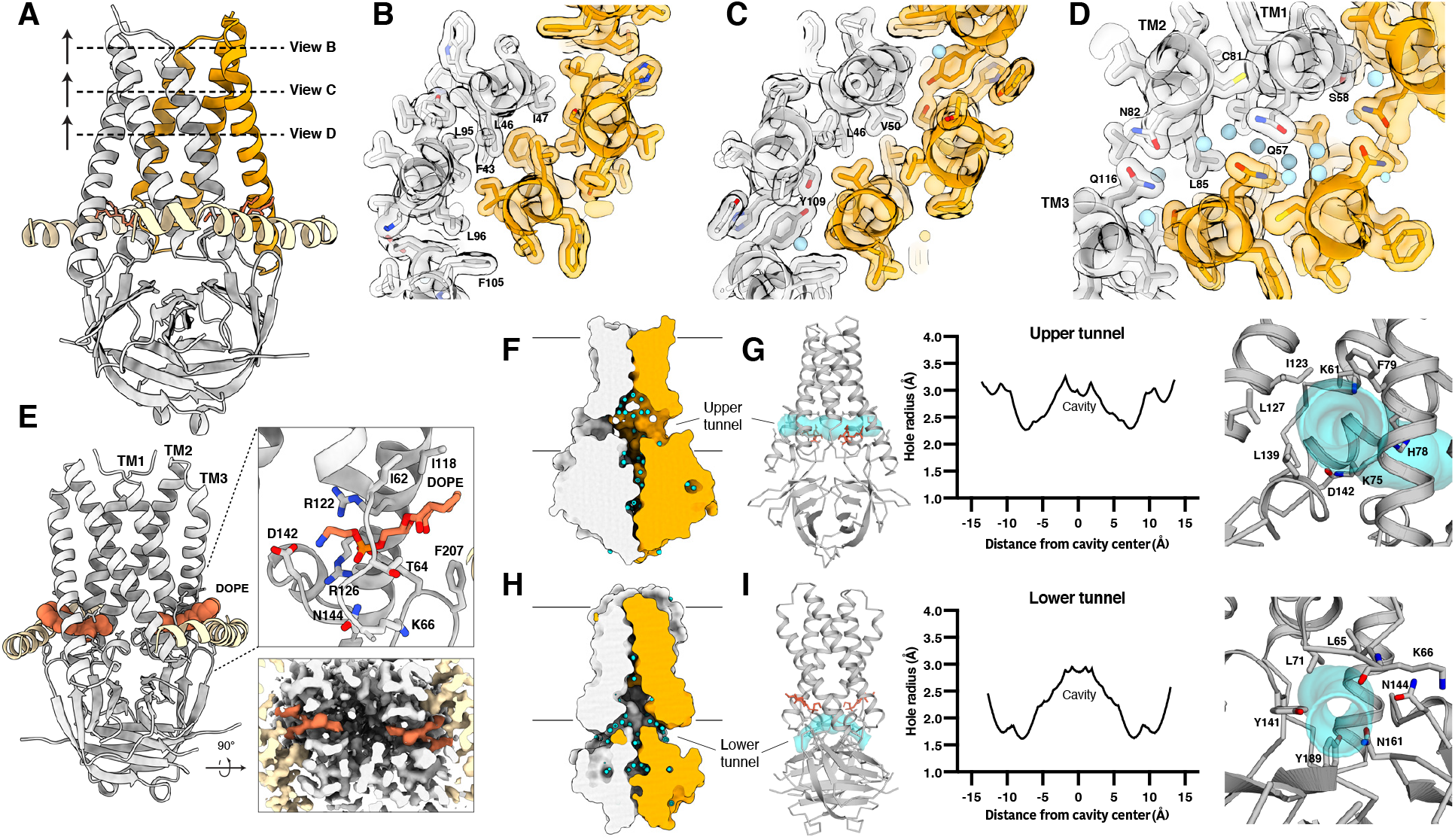
The 3a polar cavity and tunnels. (A) View of a 3a dimer from the membrane plane with planes (dotted lines) and viewing direction (arrow) for (B-D) indicated. One 3a subunit is colored orange and the other in gray. (B-D) Model and transparent surface of the viewing planes indicated in (A). Water molecules are shown as light blue spheres. (E) The intersubunit tunnel with bound DOPE lipid (coral) shown as space-filling spheres. (Right-top) Zoomed-in view of the lipid interaction with key residues shown as sticks. (Right-bottom) Cut-through view of the cryo-EM density of the lipid interaction region viewed from the cytosolic side. (F) The 3a upper tunnel with solvent-excluded surface shown for each subunit and water molecules shown as light blue spheres. The approximate lipid bilayer region is marked by black lines. (G) (Left) Model of 3a (gray) with the HOLE path through the upper tunnel shown in transparent cyan. Lipid (coral) is displayed as sticks. (Middle) Radius of the upper tunnel as a function of distance from the cavity center calculated with HOLE. (Right) View into the upper tunnel from the membrane with key residues shown as sticks. (H) As in F, for a view of the lower tunnel. (I) As in G, for the lower tunnel, with the view on the right into the tunnel from the cytosol.

While we do not observe a continuous permeation pathway across the membrane, there is a large polar cavity within the inner half of the TM region. The cavity is continuous with the cytosol and surrounding bilayer through three pairs of openings: the intersubunit, upper, and lower tunnels (Fig. 3E-I). The intersubunit tunnels run between TM1 and TM3 from opposing protomers, just above the CD, and open to the membrane-cytosol interface (Fig. 3E). Strong lipid-shaped density present within each intersubunit tunnel is modeled as DOPE (the most abundant lipid in these nanodiscs) with the ethanolamine head group pointed into the cavity. Lipid binding is stabilized by electrostatic interactions: the positively charged ethanolamine interacts with negatively charged D142 and the backbone carbonyls of I63 and D142 while the lipid phosphate interacts with R126, R122, Y206, N144, and the backbone amide of L65. The upper tunnels are formed between TM2 and TM3 within each protomer, narrow to ~2.2 Å in radius, and likely open to the membrane (Fig. 3F,G). The lower tunnels run underneath the TM1-TM2 linker and above the CD, narrow to ~2 Å in radius, and open to the cytosol (Fig. 3H,I). The lower tunnels are open paths for ion and water movement between the cell interior and channel cavity. Consistently, a network of ordered water molecules extends through these openings, the cavity, and the CD (Fig. 3F,H). Together, these features are consistent with the 3a structure representing a closed or inactivated channel.

### Biophysical properties of reconstituted 3a ion channels

We next utilized electrophysiology and calcium imaging to determine whether reconstituted 3a proteins are capable of permeating ions. We purified 3a in detergent, reconstituted it into phosphatidylcholine-containing proteoliposomes, and recorded currents across excised patches pulled from proteoliposome blisters. Excised patches from 3a-containing liposomes exhibited currents that reversed at 0 mV and displayed modest outward rectification in symmetric [K^+^], consistent with preferential sidedness of rectifying channels in the membrane after reconstitution (Fig. 4A). In contrast, channel-like activity was not observed in recordings from mock-reconstituted (empty) proteoliposomes (Fig. S15G). We evaluated ion selectivity of 3a by replacing the K^+^-containing bath solution with solutions containing Na^+^, NMDG^+^, or Ca^2+^. Solution exchange resulted in reversal potential shifts from 0.3 ± 0.3 mV in K^+^ to −8.2 ± 0.8 mV in Ca^2+^, −13.5 ± 1.8 mV in Na^+^, and −31.0 ± 1.1 mV in NMDG^+^ (Fig. 4B,D). These shifts correspond to the following permeability ratios (P_X_/P_K+_): Ca^2+^ (1.88±0.08) > K^+^ (1.0±0.01) > Na^+^ (0.59±0.04) > NMDG^+^ (0.29±0.01) (Fig. 4C). Alkaline or acidic pH had little effect on channel activity (Fig. S15F).

**Figure 4.**
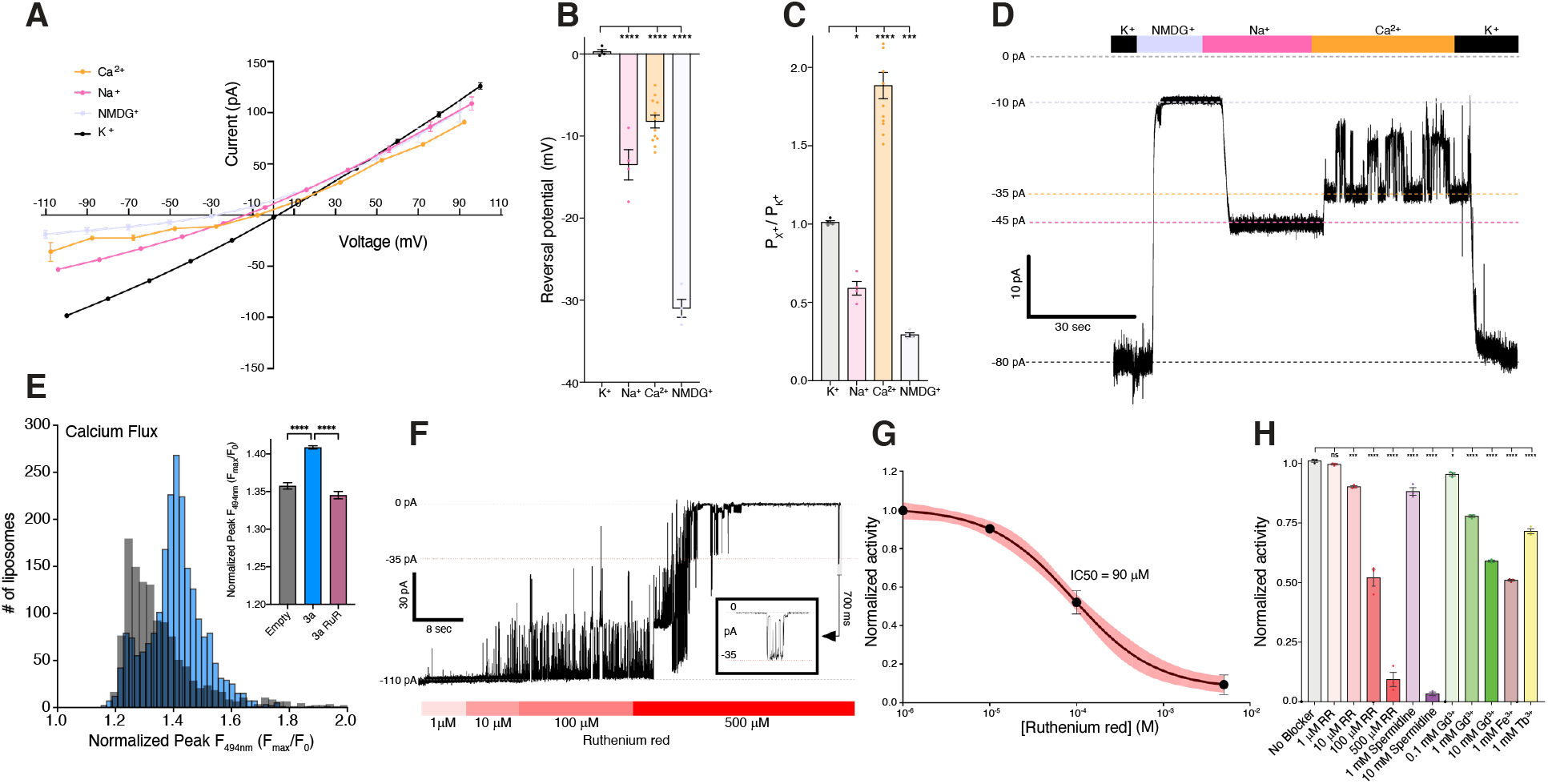
Function and inhibition of purified and reconstituted SARS-CoV-2 3a. (A) Current-voltage relationship from a 3a-proteoliposome patch. Pipette solution was 150 mM K^+^ and external solution was 150 mM K^+^ (black), 150 mM Na^+^ (pink), 75 mM Ca^2+^ (orange), or 150 mM NMDG^+^ (blue) (mean ± s.e.m., n=4-8 patches). (B) Reversal potential from (A). (C) Permeability ratios (*P_X+_* / *P_K+_*) calculated from reversal potential shifts in (B). (D) Gap-free current recording held at −80 mV during bath solution exchanges indicated in the bar above the current trace. (E) Histogram of peak calcium influx (F_max_/F) measured by Fluo-5N fluorescence following the addition of 8mM [Ca^2+^]_ext_ to 3a liposomes (blue) or empty liposomes (gray). For the inset graph, mean ± SEM of peak calcium influx for 3a liposomes, empty liposomes, and 3a liposomes incubated in 200 μM ruthenium red. (one-way ANOVA (F_(2,4945)_ = 99.01); n= 1,200-2,200 liposomes per group with Dunnett’s multiple-comparison test, **** p<0.0001). (F) Gap-free current recording in symmetric 150 mM KCl held at −80 mV during bath solution exchanges of varying ruthenium red concentrations indicated in the bar below the current trace; boxed inset: magnified channel openings and closures selected from the region indicated. (G) Normalized activity in symmetric 150 mM KCl at different concentrations of ruthenium red with fit (black line) and 95% confidence interval (red) shown (mean ± sem, n = 3 patches, IC_50_ = 90 ± 10 μM). (H) Normalized 3a activity in symmetric 150 mM KCl and (from left to right) no blocker, 1 μM ruthenium red, 10 μM ruthenium red, 100 μM ruthenium red, 500 μM ruthenium red, 1 mM spermidine, 10 mM spermidine, 0.1 mM Gd^3+^, 1 mM Gd^3+^, 10 mM Gd^3+^, 1 mM Fe^3+^, and 1 mM Tb^3+^ (mean ± sem, n = 3 patches, one-way ANOVA with Dunnett correction, *p<0.05, **p<0.01, *** p<0.001, ****p<0.0001).

Using Fluo-5N calcium imaging, we also observed significant calcium influx in 3a-proteoliposomes following the addition of 8mM [Ca^2+^]_ext_ compared to empty liposomes. Overall, these results suggest that SARS-CoV-2 3a is a Ca^2+^-permeable nonselective cation channel.

### Polycations block 3a channel activity

We next asked whether known blockers of non-selective cation channels also inhibit 3a ion conduction. We found that Ruthenium Red (RuR), a 786 Da polycationic dye, blocks 3a activity in current recordings (Fig. 4G-H) and Ca^2+^ influx measurements (Fig. 4H-I). RuR displays dose-dependent inhibition of 3a activity in proteoliposome recordings (IC50=90 ± 10 μM) with flickery block at negative potentials, similar to that observed with other large pore channels (Fig. 4G). In addition, we identified the polyamine spermidine, which showed near complete block at 10 mM, and the trivalent ions Gd^3+^, Fe^3+^, and Tb^3+^, which showed partial block at 1-10 mM, as 3a inhibitors in proteoliposome recordings (Fig. 4G). RuR, spermidine, Gd^3+^, and other trivalents are relatively nonselective inhibitors of cation channels including TRPs, RyRs, CALHMs, K2Ps, and KIRs^29–35^. SARS-CoV-2 3a activity was not inhibited by Ba^2+^ or the small molecule emodin (Fig. S15D,E), two reported inhibitors of SARS-CoV-1 3a^23,24^. This is consistent with our finding that the addition of emodin had no effect on SARS-CoV-2 3a structure (Fig. S7-9).

### 3a permeates the large divalent ion YO-PRO-1

Given the ability of 3a channels to conduct NMDG^+^ (Fig. 4A-D), we next used a YO-PRO-1 fluorescence-based flux assay to assess the ability of 3a channels to conduct other large cations. YO-PRO-1, a 629 Da divalent cation, does not readily cross lipid bilayers and has been used to study the activity of other non-selective cation channels, including P2X7 and TRP channels, that conduct large organic cations^29,30,36,37^. Robust YO-PRO-1 uptake was observed in a subset of 3a liposomes that was significantly higher than background fluorescence observed in empty liposomes (Fig. 5A-D). To compare YO-PRO-1 uptake across multiple experiments, we quantified a single value (area under the curve) to represent both the number of YO-PRO-1+ liposomes and the amount of YO-PRO-1 uptake (Fig. 5E). Using this analysis, we found 3a-liposomes display significantly greater YO-PRO-1 uptake and accumulation as compared to empty liposomes (Fig. 5E). Uptake was dependent on 3a protein because liposomes containing the human K2P channel TRAAK displayed little uptake and accumulation of YO-PRO-1, similar to empty liposomes, as expected for a highly selective K+ channel (Fig. 5A, C-E). Similar to current recordings (Fig. 4E-F), we observed a dose-dependent RuR block of 3a activity in the YO-PRO-1 flux assay (IC50=175 ± 98 μM). These data demonstrate that 3a can form an ion channel that is capable of passing large cations. RuR (786 Da) block and YO-PRO-1 (629 Da) and NMDG^+^ (406 Da) permeation sets limits on 3a pore size that are similar those observed for TRP and P2X channels.

**Figure 5.**
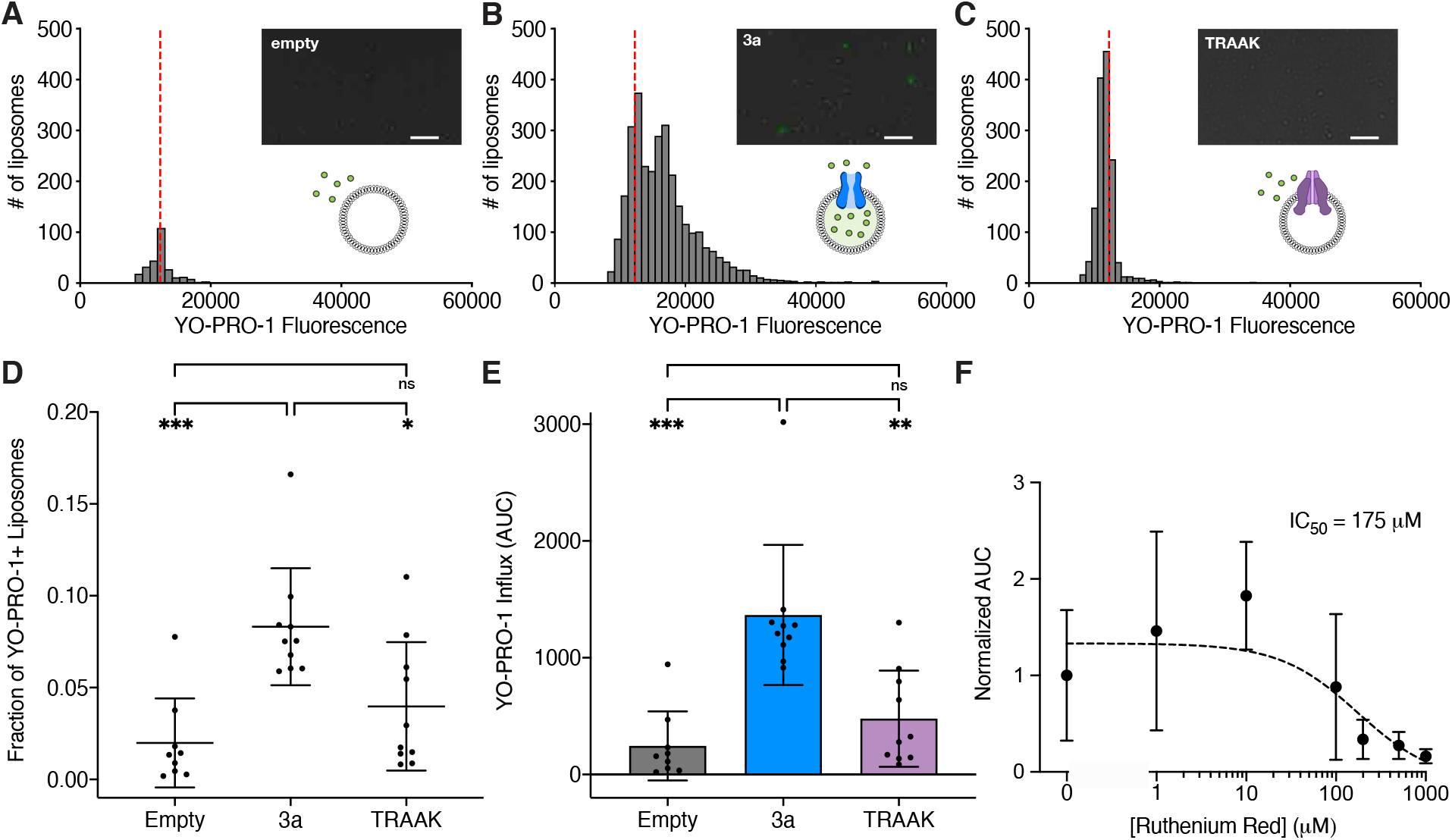
A flux assay of SARS-CoV-2 3a channel activity. (A-C) Histograms from quantified images following a 10-minute incubation in 10 μM YO-PRO-1 for (A) empty liposomes, (B) 3a liposomes, or (C) TRAAK liposomes (n = 255, 3046, and 1408 for empty, 3a, and TRAAK liposomes). Dotted red line indicates the position of the mean fluorescence of empty liposomes. (Insets) Representative images with YO-PRO-1 fluorescence (green) overlaid on brightfield. Scale bar = 20μm. (D) Fraction of YO-PRO-1-positive liposomes (mean ± SD, n = 9-10 independent replicates with 500-5,000 liposomes per replicate, *p* = 0.0004 (one-way ANOVA (F_(2,26)_ = 10.52)), Tukey’s multiple-comparison test: *p*_empty vs. 3a_ = 0.0004; *p*_TRAAK vs. 3a_ = 0.0113; *p*_empty vs. TRAAK_ = 0.3571). (E) Area under the curve (AUC) of YO-PRO-1 fluorescence histograms of the fraction of liposomes that took up YO-PRO-1 per well (mean ± SD, n= 9-10 independent wells per group of ~500-5,000 liposomes per well, *p* < 0.0001 (Welch’s ANOVA test (W_(2,16.68)_ =13.22)), Dunnett’s multiple-comparison test: *p*_empty vs. 3a_ =0.0005; *p*_TRAAK vs. 3a_ = 0.0041; *p*_empty vs. TRAAK_ = 0.4178). (F) YO-PRO-1 uptake in 3a-liposomes is inhibited by ruthenium red. Sigmoidal fit (black dotted line) and IC_50_ are shown.

### 3a mutants alter channel activity

We speculate that an open conformation of 3a involves substantial rearrangement of the hydrophobic seal to create a central pore through the membrane or more subtle rearrangement of the partially hydrophilic membrane facing sides of TM2 and TM3 to create membrane facing conduction pathways. To probe the structural determinants for 3a ion conduction, we set out to identify 3a mutations that alter channel activity. To this end, we purified nine mutated or truncated 3a constructs, reconstituted each into proteoliposomes, and compared them to wild-type 3a in electrophysiological recordings (Fig. 6A, S16). Consistent with our hypothesis that 3a channel opening would likely require conformational changes in transmembrane helices to expand the constrictions above the polar cavity to the extracellular side, we find that two separate mutants at the top of the cavity (Q57E and S58L/Q116L) alter channel activity. Both Q57E and S58L/Q116L mutants reduce Ca^2+^ and NMDG^+^ permeability, but did not alter Na^+^ or K^+^ permeability. These effects were specific to these mutations as seven additional mutations, including a common circulating variant Q57H^38^, had no effect on relative ion permeability.

**Figure 6.**
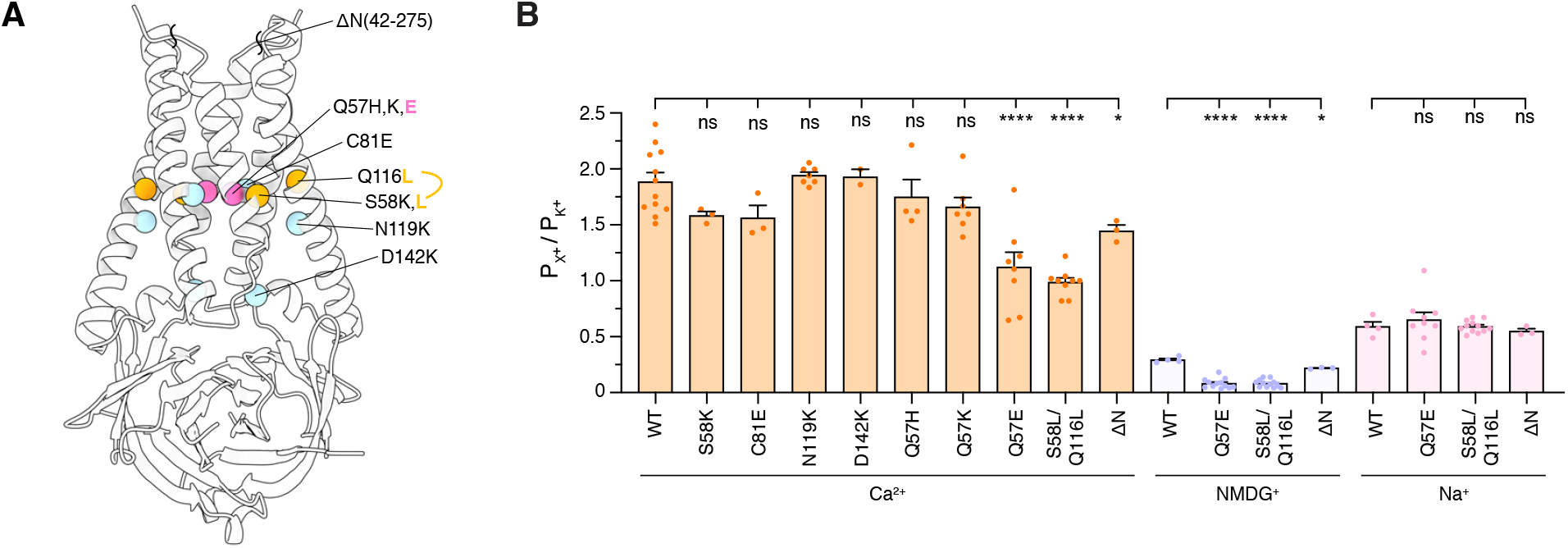
Modification of 3a alters channel activity. (A) View of 3a from the membrane plane with the positions of 3a mutations and truncation indicated. (B) Permeability ratios (*P_X_* / *P_K+_*) calculated from reversal potential shifts for 3a mutations and ΔN truncation (mean ± s.e.m., n=8, 3, 3, 7, 2, 4, 7, 8, 9, and 3 patches for wild-type, S58K, C81E, N119K, D142K, Q57H, Q57K, Q57E, S58L/Q116L, and ΔN, respectively, *p<0.05, **** p<0.0001, one-way ANOVA with Dunnett correction for mltiple comparisons).

Our structural analysis revealed an unassigned region of density in the cryo-EM maps stretching between subunits just above the extracellular side of the transmembrane region that likely corresponds to a portion of the unmodeled N-terminus (Fig. S14). To investigate the potential role of the N-terminus in channel function, we generated an N-terminal deletion construct lacking the first 41 amino acids (3aΔN). This mutant displayed reduced Ca^2+^ and NMDG^+^ relative permeability in proteoliposome recordings, albeit to a lesser extent than Q57E and S58L/Q116L (Figure 6B). Changes in permeability may reflect changes in pore properties or altered cellular localization. The relative permeability changes observed in liposomes combined with the observation that 3aΔN channels localize to the plasma membrane (Fig. S14) suggest that the N-terminus mediates the ability of 3a to permeate large cations.

### 3a-like proteins in *Alpha- and Beta-coronavirus*

While 3a is well conserved in the *Betacoronavirus* subgenus (Fig. S1), related proteins have not yet been identified in other coronaviruses, including the other five species known to infect humans: MERS-CoV, HCoV-NL63, HCoV-229E, HCoV-HKU1, and HCoV-OC43. Thus, we asked whether we could identify more distant homologs using the 3a structure and structure prediction algorithms. 3a homologs were not detected in *Gammacoronavirus*, *Deltacoronavirus*, or in the *Betacoronavirus* subgenus *Embecovirus* (which includes HCoV-HKU1 and HCoV-OC43). Distant homology to the CD was identified in the membrane protein ORF5 found in *Betacoronavirus* subgenus *Merbecovirus* species including MERS-CoV. However, we identified high confidence structural homologs in all remaining *Betacoronavirus* and *Alphacoronavirus* subgenera (including in HCoV-229E and HCoV-NL63 (Fig. S17)), several of which have been reported to have ion channel activity^39–41^. Notably, coronaviruses with 3a homologs have been proposed to derive from viruses that circulate primarily in bats, while coronaviruses without 3a structural homologs have been proposed to derive from viruses that circulate primarily in rodents, birds, or pigs (Table 2). Perhaps the presence of 3a in coronaviruses with bats as their principal reservoir reflects a unique aspect of bat coronavirus biology.

Overall, we demonstrate that SARS-CoV-2 3a forms a non-selective Ca^2+^ permeable cation channel when reconstituted in liposomes. 3a structures show a novel fold with a polar cytoplasmic cavity open to the cytosol and membrane below a hydrophobic seal, consistent with a closed or inhibited state. The basis for 3a channel gating and conduction remains to be determined. SARS-CoV-2 3a channels display permeability to large cations including NMDG^+^ and YO-PRO-1 and sensitivity to RuR and Gd^3+^, reminiscent of other Ca^2+^-permeable channels including TRPV1, TRPA1, and P2X7^29,30^. Channel activity may be important for promoting viral maturation in cells through inhibition of autophagy and disruption of lysosomes^7,8^. The calcium permeability of 3a is of particular interest in the context of infection. SARS-CoV-2 3a has been recently shown to trigger programmed cell death *in vitro*^26^ and calcium-influx through 3a may be the switch that activates calcium-dependent caspases and apoptosis. Previous studies showed SARS-CoV-1 3a channel expression in infected lung pneumocytes, a key cell type also infected by SARS-CoV-2, and calcium signaling in type II pneumocytes plays an important role in maintaining airway homeostasis^42,43^. Thus, the expression of a calcium permeable channel, like SARS-CoV-2 3a, could impact lung homeostasis and COVID-19 pathogenesis. Our data suggest that 3a may represent a new target for treating COVID-19 and other coronavirus diseases and points the way to future experiments that will elucidate the role of 3a in the viral life cycle and disease pathology.

## Methods

### Cloning and protein expression

The coding sequence for the 3a protein from SARS-Cov-2 was codon optimized for *Spodoptera frugiperda* (Sf9 cells) and synthesized (IDT, Newark, NJ). The sequence was then cloned into a custom vector based on the pACEBAC1 backbone (MultiBac; Geneva Biotech, Geneva, Switzerland) with an added C-terminal PreScission protease (PPX) cleavage site, linker sequence, superfolder GFP (sfGFP) and 7xHis tag, generating a construct for expression of 3a-SNS-LEVLFQGP-SRGGSGAAAGSGSGS-sfGFP-GSS-7xHis. Mutants and truncation were also introduced into this construct using PCR. MultiBac cells were used to generate a Bacmid according to manufacturer’s instructions. Sf9 cells were cultured in ESF 921 medium (Expression Systems, Davis, CA) and P1 virus was generated from cells transfected with Escort IV reagent (MillaporeSigma, Burlington, MA) according to manufacturer’s instructions. P2 virus was then generated by infecting cells at 2 million cells/mL with P1 virus at a MOI ~0.1, with infection monitored by fluorescence and harvested at 72 hours. P3 virus was generated in a similar manner to expand the viral stock. The P2 or P3 viral stock was then used to infect Sf9 cells at 4 million cells/mL at a MOI ~2–5. At 72 hours, infected cells containing expressed 3a-sfGFP protein were harvested by centrifugation at 2500 x g for 10 minutes and frozen at −80°C.

### Protein purification

For preparation of the 3a dimer and mutant constructs, infected Sf9 cells from 1 L of culture (~15-20 mL of cell pellet) were thawed in 100 mL of Lysis Buffer containing 50 mM Tris, 150 mM KCl, 1mM EDTA pH 8. Protease inhibitors (Final Concentrations: E64 (1 μM), Pepstatin A (1 μg/mL), Soy Trypsin Inhibitor (10 μg/mL), Benzamidine (1 mM), Aprotinin (1 μg/mL), Leupeptin (1μg/mL), AEBSF (1mM), and PMSF (1mM)) were added to the lysis buffer immediately before use. Benzonase (4 μl) was added after the cell pellet thawed. Cells were then lysed by sonication and centrifuged at 150,000 x g for 45 minutes. The supernatant was discarded and residual nucleic acid was removed from the top of the membrane pellet using DPBS. Membrane pellets were scooped into a dounce homogenizer containing Extraction Buffer (50 mM Tris, 150 mM KCl, 1 mM EDTA, 1% n-Dodecyl-β-D-Maltopyranoside (DDM, Anatrace, Maumee, OH), pH 8). A 10% stock solution of DDM was dissolved and clarified by bath sonication in 200 mM Tris pH 8 prior to addition to buffer to the indicated final concentration. Membrane pellets were then homogenized in Extraction Buffer and this mixture (150 mL final volume) was gently stirred at 4°C for 1 hour. The extraction mixture was centrifuged at 33,000 x g for 45 minutes and the supernatant, containing solubilized membrane protein, was bound to 4 mL of Sepharose resin coupled to anti-GFP nanobody for 1 hour at 4°C. The resin was then collected in a column and washed with 10 mL of Buffer 1 (20 mM HEPES, 150 mM KCl, 1 mM EDTA, 0.025% DDM, pH 7.4), 40 mL of Buffer 2 (20 mM HEPES, 500 mM KCl, 1 mM EDTA, 0.025% DDM, pH 7.4), and 10 mL of Buffer 1. The resin was then resuspended in 6 mL of Buffer 1 with 0.5 mg of PPX protease and rocked gently in the capped column for 2 hours. Cleaved 3a protein was then eluted with an additional 8 mL of Wash Buffer, spin concentrated to ~500 μl with Amicon Ultra spin concentrator 10 kDa cutoff (Millipore), and then loaded onto a Superdex 200 increase column (GE Healthcare, Chicago, IL) on an NGC system (Bio-Rad, Hercules, CA) equilibrated in Buffer 1. Peak fractions containing 3a channel were then collected and spin concentrated prior to incorporation into proteoliposomes or nanodiscs. For the tetramer, the preparation was carried out in a similar manner, except with overnight protease cleavage and collection of a peak of larger hydrodynamic radius (see Fig. S3).

### Proteoliposome formation

For proteoliposome patching experiments, we incorporated protein into lipid and generated proteoliposome blisters for patch recordings using dehydration and rehydration as described previously with the following modifications^44^. 3a dimer was first purified into Buffer 1. Protein was then exchanged into lipid with the addition of Biobeads SM2 (Bio-Rad, Hercules, CA) and an overnight incubation at a protein:lipid ratio of 1:10 (corresponding to 0.5 mg purified 3a dimer and 5 mg of cleared Soybean L-α-phosphatidylcholine (Soy PC, MillaporeSigma, Burlington, MA) in DR Buffer (5 mM HEPES, 200 mM KCl, pH7.2). For the YO-PRO-1 assay, 3a was incorporated at a ratio of 1:50. TRAAK control proteoliposomes were prepared at 1:50 as described previously^44^. Control liposomes were prepared from the same lipid mix and protocol with protein replaced with Buffer 1.

### Electrophysiology

All electrophysiology recordings were made from 3a-reconstituted Soy PC proteoliposomes. Patches formed in an inside-out configuration and were quickly (within 5-10 seconds) transferred to a solution exchange chamber. Recordings were made at room temperature using Clampex 10.7 data acquisition software with an Axopatch 200B Patch Clamp amplifier and Digidata 1550B digitizer (Molecular Devices) at a bandwidth of 1 kHz and digitized at 500 kHz. A pressure clamp (ALA Scientific) was used to form seals. Potassium pipette and bath solution was 5 mM HEPES pH 7.2, 150 mM KCl. Sodium bath solution was 5 mM HEPES pH 7.2, 150 mM NaCl. NaCl in the bath solution was substituted for 150 mM NMDG-Cl or 75 mM CaCl_2_ for permeability experiments. Borosilicate glass pipettes were pulled and polished to a resistance of 2-5 MΩ when filled with pipette solution. For cation permeability experiments, liquid junction potentials were calculated and data were corrected offline. For current-voltage plots, the following voltage protocol was applied: V_hold_= 0 mV mV; V_test_= −100 to +100 mV, Δ20 mV, t_test_ = 1 second. Currents from each patch correspond to mean values during the step to the indicated voltage.

Permeability ratios were calculated according to Goldman-Hodgkin-Katz relationship. For monovalent cations, permeability ratios were calculated as P_X+_/P_K+_ = exp(ΔV_rev_F/RT). For divalent cations, permeability ratios were calculated as: P_X2+_/P_K+_ = *α*_K+_[K+]exp(ΔV_rev_F/RT)(1 + exp(ΔV_rev_F/RT))/ 4*α*_X2+_[X^2+^] where V_rev_ is the reversal potential, F is Faraday’s constant, R is the universal gas constant, and T is absolute temperature (where RT/F = 25.2 mV at room temperature), and *α* is the ion activity coefficient (assumed to be 0.75 for K^+^ and 0.25 for Ca^2+^).

### Ca^2+^ uptake assay in proteoliposomes

Fluo-5N was incorporated into 3a-reconstituted proteoliposomes and control liposomes by first thawing frozen liposomes at a ratio of ~1:20 (v:v) into modified sucrose formation buffer^45^ 25 mM HEPES pH 7.4, 150 mM KCl, and 262 mM Sucrose with a final concentration of 25 uM Fluo-5N (Stock concentration 5 mM in DMSO, AAT Bioquest). Next, the liposome mixtures in Eppendorf tubes were placed in a foam flotation in an ice bath and sonicated with a Branson Digital Sonifier 450 for 30 seconds total (10% Amplitude, 10 second pulse with 59 second wait time). Excess (unincorporated) Fluo-5N was then removed using microspin G-50 columns (Cytiva). The sample was then diluted into solution containing 50mM HEPES, 300mM KCl, and 2mM EGTA, pH=7.4 and plated onto poly-D-lysine (Sigma-Aldrich, 1mg/ml) coated 96-well plates, and centrifuged (5,000 x g for 5 min) at room temperature. Ca^2+^ influx was measured upon addition of a 50 mM HEPES, 285 mM KCl, 10 mM CaCl_2_, pH=7.4 solution. Images were acquired every 3 seconds for a total of 150 seconds. Fluorescence intensity of each liposome was normalized to its average fluorescence prior to Ca2+ addition.

### YO-PRO-1 flux assay

Liposomes were diluted in saline solution containing 300mM KCl, 50mM HEPES, pH=7.4 and plated onto poly-D-lysine (Sigma-Aldrich, 1mg/ml) coated 96-well plates, centrifuged (5,000 x g for 5 min) and incubated in 10μM YO-PRO-1 iodide (Invitrogen) for 10 min at room temperature. Liposomes were rinsed with saline solution to remove free YO-PRO-1. Images were acquired using an ImageXpress Micro XLS microscope with a solid-state light source and 20X air objective. Images were analyzed using MetaXpress 6 software (Molecular Devices). Empty liposomes displayed low-level fluorescence in the 488/540nm range. Thus, liposomes were defined as YO-PRO-1+ if their fluorescence intensity was above the average intensity of unlabeled, empty liposomes. To calculate the total number of liposomes, we quantified the total number of liposomes using auto segmentation on MetaXpress 6 from brightfield images. Stock solutions of Ruthenium red (Tocris) were prepared in water, diluted in bath solution and applied to liposomes or cells 10min before the start of the experiment. All statistical tests were performed using Prism (GraphPad). Values are reported as the mean ± SD or mean ± SEM as indicated. Mann Whitney or a one-way or Welch’s ANOVA followed by the Dunnett’s, Sidak’s, or Tukey’s post hoc tests (where appropriate) were used for statistical comparisons.

### Nanodisc Formation

Freshly purified 3a dimer in Buffer 1 was reconstituted into MSP1E3D1 nanodiscs with a mixture of lipids (DOPE:POPS:POPC at a 2:1:1 mass ratio, Avanti, Alabaster, Alabama) at a final molar ratio of 1:4:400 (Monomer Ratio: 3a, MSP1E3D1, lipid mixture). First, 20 mM solubilized lipid in Nanodisc Formation Buffer (20 mM HEPES, 150 mM KCl, 1 mM EDTA pH 7.4) was mixed with additional DDM detergent and 3a protein. This solution was mixed at 4°C for 30 minutes before addition of purified MSP1E3D1. This addition brought the final concentrations to approximately 15 μM 3a, 60 μM MSP1E3D1, 6 mM lipid mix, and 10 mM DDM in Nanodisc Formation Buffer. The solution with MSP1E3D1 was mixed at 4°C for 10 minutes before addition of 200 mg of Biobeads SM2. Biobeads (washed into methanol, water, and then Nanodisc Formation Buffer) were weighed after liquid was removed by pipetting (damp weight). This mix was incubated at 4°C for 30 minutes before addition of another 200 mg of Biobeads (for a total 400 mg of Biobeads per 0.5 mL reaction). This final mixture was then gently tumbled at 4°C overnight (~ 12 hours). Supernatant was cleared of beads by letting large beads settle and carefully removing liquid with a pipette. Sample was spun for 10 minutes at 21,000 x g before loading onto a Superdex 200 increase column in 20 mM HEPES, 150 mM KCl, pH 7.4. Peak fractions corresponding to 3a protein in MSP1E3D1 were collected, 10 kDa cutoff spin concentrated and used for grid preparation. MSP1E3D1 was prepared as described^46^ without cleavage of the His-tag. Tetrameric 3a in nanodiscs was prepared similarly, except with a ratio of 1:2:200 (Monomer Ratio: 3a, MSP1E3D1, lipid mixture).

### Grid Preparation

Dimeric 3a in MSP1E3D1 was prepared at final concentration of 1.1 mg/mL. For the sample with emodin (MillaporeSigma, Burlington, MA, Catalog# E7881), a stock solution of 50 mM emodin in DMSO added to protein sample for final concentrations of 1.1 mg/mL 3a and 100 μM emodin and 1% DMSO. Concentrated sample was cleared by a 10 minute 21,000 x g spin at 4°C prior to grid making. For freezing grids, a 3 μl drop of protein was applied to freshly glow discharged Holey Carbon, 300 mesh R 1.2/1.3 gold grids (Quantifoil, Großlöbichau, Germany). A FEI Vitrobot Mark IV (ThermoFisher Scientific) was used with 4°C, 100% humidity, 1 blot force, a wait time of ~5 seconds, and a 3 second blot time, before plunge freezing in liquid ethane. Grids were then clipped and used for data collection. Tetrameric 3a in MSP1E3D1 was frozen at 0.7 mg/mL with the same grid preparation. Unimaged grids from the same session were then shipped to the Netherlands for data collection on the 300 kV microscope with Falcon 4/CFEG/Selectris energy filter.

### Cryo-EM data acquisition

For data collected on the Talos Arctica, grids were clipped and transferred to the microscope operated at 200 kV. Fifty frame movies were recorded on a Gatan K3 Summit direct electron detector in super-resolution counting mode with pixel size of 0.5685 Å. For the apo 3a dataset, the electron dose was 9.528 e^−^ Å^2^ s^−1^ and 10.135 e^−^ Å^2^ s^−^ and total dose was 50.02 e^−^ Å^2^ and 53.72 e^−^ Å^2^ in the first set (1-2007) and second set (2008-6309) of movies respectively. The two different doses are the result of needing to restart the electron gun during collection. For the 3a with added emodin dataset, the electron dose was 8.991 e^−^ Å^2^ s^−1^ and total dose was 47.21 e^−^ Å^2^. For the 3a tetramer, the electron dose was 8.841 e^−^ Å^2^ s^−1^ and total dose was 49.95 e^−^ Å^2^. Nine movies were collected around a central hole position with image shift and defocus was varied from −0.6 to −2.0 μm through SerialEM^47^.

For data collected for the high resolution 3a dimer structure, clipped grids from the same batch used on Talos Artica were sent to Thermo Fisher Scientific RnD division in Eindhoven, The Netherlands. Grids were loaded onto the Krios G4 microscope equipped Cold Field Emission gun (CFEG) operated at 300 kV. Data were collected on a Falcon 4 detector that was mounted behind a Selectris X energy filter. The slit width of the energy filter was set to 10eV. 5599 movie stacks containing 1429 raw frames were collected with EER (electron event representation) mode^48^ of Falcon 4 detector at a magnification of 165 kX corresponding to a pixel size of 0.727 Å. Each movie stack was recorded with an exposure time of 6s with a total dose of 50e/A2 on sample and a defocus range between 0.5 to 1.2 μm.

See Table 1 for detailed data collection statistics.

### Cryo-EM data processing

For the apo 3a dimer, motion-correction and dose-weighting were performed on all 6,309 movies using RELION 3.1’s implementation of MotionCor2, and 2x “binned” to 1.137 Å per pixel^49–51^. CTFFIND-4.1 was used to estimate the contrast transfer function (CTF) parameters^52^. Micrographs were then manually sorted to eliminate subjectively bad micrographs, such as empty or contaminated holes, resulting in 3,611 good micrographs. Additionally, micrographs with a CTF maximum resolution lower than 4 Å were discarded, resulting in 2,595 remaining micrographs. Template-free auto-picking of particles was performed with RELION3.1’s Laplacian-of-Gaussian (LoG) filter yielding an initial set of particles. This initial set of particles were iteratively classified to generate templates, which were subsequently used to template-based auto-pick 1,750,730 particles.

Template picked particles were iteratively 2D-classified in RELION3.1 and then in cryoSPARC v2^53^, resulting in 820,543 particles. These particles were subsequently 3D-classified in cryoSPARC v2 with iterative ab-initio and heterogeneous refinement jobs. The resulting maps were visually evaluated with regard to the transmembrane domain density. A set of 86,479 particles were identified, polished in RELION3.1 and refined in cryoSPARC v2 with subsequent homogeneous and non-uniform refinement^54^ jobs (maps were low-pass filtered to an initial resolution where TM density was still visible (6-9 Å), and the dynamic mask was tightened with the near (2-5 Å) and far (3-9 Å) parameters), yielding a map with overall resolution of 3.6 Å. UCSF pyem tools were used to convert data from cryoSPARC to RELION format^55^.

From this set of 86,479 particles, 2D-classification was performed in RELION3.1 to identify a set of particles with subjectively equal view distribution. From the resulting set, 1,000 particles were randomly sampled and their coordinates used for training in the Topaz particle-picking pipeline^56^. Training, picking, and extraction were performed independently on each subset of the micrographs. 4,134,279 total particles were extracted in RELION3.1 with a box size of 256 pixels and “binned” 4x to 4.548 Å/pixel. These particles were then iteratively 2D-classified in RELION3.1 resulting in 2,674,606 particles which were extracted at 2.274 Å/pixel. 2D-classification was continued in both RELION3.1 and cryoSPARC v2 resulting in 1,429,763 particles. Further classification was performed in cryoSPARC v2 with subsequent ab-initio (4 classes, max resolution 8 Å) and heterogeneous refinement (8 Å initial resolution) jobs. The two best classes were selected and the particles pooled resulting in 743,800 particles which were extracted in RELION3.1 at 1.137 Å/pixel.

Iterative 3D-classification was performed with subsequent ab-initio and heterogeneous refinement jobs as described above. Following each round, 2D classification jobs were used to “rescue” good particles from the worst classes before the next round. After 3 rounds, a final 2D-classificaiton job was used to identify 112,502 particles, which were subsequently pooled with the previous 86,479 RELION3.1 template-picked particles, resulting in 185,871 particles after duplicates (within 100 Å) were removed with RELION3.1.

These particles were then refined with subsequent homogeneous and non-uniform refinement jobs resulting in a map with overall resolution of 3.4 Å. This map was post-processed in RELION3.1 using a mask with a soft edge (5 pixel extension, 7 pixel soft-edge), the output of which was used for Bayesian particle polishing in RELION3.1 (training and polishing were each performed independently on each subset of the micrographs). The resulting “shiny” particles were then refined in cryoSPARC v2 with subsequent homogenous refinement (1 extra pass, 7 Å initial resolution) and non-uniform refinement (C2, 1 extra pass, 9 Å initial resolution) to yield a map with 2.9 Å overall resolution.

For 3a dimer with added 100 μM emodin, initial processing was similar to the dimer without added drug (see Fig. S8). As with the apo 3a dimer, the critical steps included Topaz particle picking, particle clean-up with cryoSPARC v2 ab-initio and heterogeneous refinement, non-uniform refinement with tightened masking, and RELION3.1 Bayesian particle polishing. However, in contrast to the apo dataset, we observed a set of particles that were included in < 4 Å reconstructions that had discontinuous transmembrane domain density. Removal of these particles with RELION3.1 3D classification without angular sampling led to the best map from the emodin-added dataset. We did not see any evidence of bound emodin, but the 1% DMSO added with drug addition may have contributed to subtle map differences (Fig. S8,S9).

For the 3a tetramer, the initial 7,092 micrographs were first cleaned using manual inspection and removal of images with < 4 Å CtfMaxResolution to obtain a set of 4,324 micrographs. Reference particles for Topaz particle picking were generated by first template picking in RELION3.1, followed by 2D classification in both RELION3.1 and cryoSPARC v2, and subsequent ab-initio in cryoSPARC v2. Particles from various views were then selected from iterative RELION3.1 2D classification to create a set of 6,843 particles. Using these coordinates for training, Topaz particle picking was then performed to generate a set of 1,282,913 initial particles. These particles were then cleaned using 2D classification in RELION3.1 and cryoSPARC v2, followed by rounds of cryoSPARC v2 ab-initio and RELION3.1 3D classification. A major hurdle for tetramer processing was obtaining a reconstruction where most particles were properly oriented in the same direction (i.e. ICD domains on the same side of the nanodisc as seen in the 2D classes, see Fig. 2F).

Substantial, cleanup by 3D-classification was needed to generate a correctly aligned reference map, but this map could then be used as a reference for refinements and classification for larger particle sets. Reconstructions with C1 or C2 symmetry looked similar (see Fig. S10), although no tetramer reconstruction went to high enough resolution to determine symmetry with certainty. Therefore, it is possible that either the tetramer is pseudosymmetric or that different particles have heterogeneous orientations between dimer pairs. For the tetramer, the highest resolution reconstruction came from cryoSPARC v2 non-uniform refinement with a tightened mask, which was subsequently used for dimer-docking and figure preparation.

For the apo 3a dimer imaged on a Krios with CFEG, Selectris X, and Falcon 4, the EER movie motion correction and subsequent polishing was performed in RELION3.1 using the devel-eer branch of Relion. Dose fractions consisting of 30 frames corresponding to 1.035e/A2 per fraction were created. The initial 5599 micrographs were pruned by selecting 4495 micrographs with < 3.5 Å CtfMaxResolution that also passed manual inspection. Similarly to the Arctica datasets, an initial set of particles was generated with template based picking and subsequent 2D classification, Ab initio, and Heterogeneous refinement for clean-up was performed in cryoSPARC v2 (See Fig. S11). This particle set (44,944) was then used as training for the first round of Topaz particle picking. For initial particle clean-up, the most important step was heterogeneous refinement in cryoSPARC v2, yielding a particle set (215,227) that gave a C2 non-uniform refinement at 2.69 Å. For final particle clean-up, using RELION3.1, polished particles were subjected to 3D classification with no angular sampling and high tau (40) during which we monitored the rounds of classification by looking for convergence in resolution and inspecting maps to check for quality of protein features or noise. This type of job reliably allowed us to select the best particles (61,531) for high-resolution reconstruction at 2.26 Å.

At this point, a second Topaz training and picking round was conducted using this set of particles. After merging the best particles from both rounds of Topaz picking, removing potential duplicate particles within a distance of 100 Å, and subsequent processing, we obtained a set of particles (91,218) that achieved a non-uniform reconstruction at 2.17 Å. Finally, rounds of CtfRefine in cryoSPARC v2 and a final non-uniform refinement (with higher-order CTF terms enabled) achieved a map with an estimated resolution of 2.1 Å. Depending on inputs into sharpening and local resolution jobs (Fig. S12A), the map has regions that give resolution estimates significantly below 2 Å, consistent with our observations of holes in aromatic side chain density.

We note that the merging of two Topaz picked particle sets likely allowed us to break a particle-limited barrier to achieve the final reconstruction based on ResLog^57^ analysis performed in cryoSPARC (See Fig. S12C). Finally, we sampled various box sizes during processing (as large as 416 pixels and as small as 256 pixels)^58^. The best reconstructions were consistently achieved with our final box size of 300 pixels (218.1 Å).

### Modeling, Refinement, and Analysis

Apo dimeric 3a cryo-EM maps were sharpened using cryoSPARC and were of sufficient quality for de novo model building in Coot^59^. Real space refinement of the models was carried out using Phenix.real_space_refine^60^. Molprobity^61^ was used to evaluate the stereochemistry and geometry of the structure for subsequent rounds of manual adjustment in Coot and refinement in Phenix. For final sharpening and visualization of the high-resolution map we used Phenix Resolve Density Modification^62^. Docking of the apo dimeric 3a into the tetrameric 3a cryo-EM map was performed in Phenix using a map in which large empty regions of the nanodisc were erased in Chimera^63^. Similar results were found using maps with only the CDs present. Cavity measurements were made with HOLE^64^ implemented in Coot. Comparisons to the structure database was performed with DALI^28^. Structure prediction was performed with Phyre2^65^. Figures were prepared using PyMOL, Chimera, ChimeraX^66^, Fiji, Prism, GNU Image Manipulation Program, and Adobe Photoshop and Illustrator software.

### Fluorescence Size Exclusion Chromatography (FSEC)

Sf9 cells (~4 million) from the third day of infection were pelleted, frozen, and then thawed into extraction buffer (20mM Tris pH 8, 150 mM KCl, all protease inhibitors used for protein purification, 1 mM EDTA, 1% DDM). Extraction was performed at 4°C for 1 hour and lysate was then pelleted at 21,000 x g at 4°C for 1 hour to clear supernatant. Supernatant was then run on a Superose 6 Increase column with fluorescence detection for GFP into 20 mM HEPES pH 7.4, 150 mM KCl, .025% DDM.

### Transfection for Confocal Imaging

The constructs for full length 3a and 3aΔN were cloned into a vector with a CMV-promoter and C-terminal EGFP. Constructs (2 μg) were transfected into HEK293 cells on glass coverslips using Fugene HD (Promega, Madison, WI) per manufacturer’s instructions. Two days after transfection cells were washed with DPBS and then fixed in 4% Formaldehyde in DPBS for 10 minutes. Cells were then washed with DPBS before mounting the coverslip with Prolong Glass Antifade with NucBlue (ThermoFisher Scientific) per manufacturer’s instructions. Fluorescent images were collected using a Zeiss LSM 880 NLO AxioExaminer confocal microscope at either 20X (NA 1.0) or 63X oil immersion objective (NA 1.4). The samples were excited with 488nm argon laser and image analysis was performed using ImageJ.

## Data and reagent availability

All data and reagents associated with this study are publicly available. For dimeric apo 3a, the final model is in the PDB under 6XDC, the final map is in the EMDB under EMD-22136, and the original micrograph movies and final particle stack is in EMPIAR under EMPIAR-10439. For tetrameric apo 3a, the final map is in the EMDB under EMD-22138 and the original micrograph movies and final particle stack is in EMPIAR under EMPIAR-10441. For dimeric 3a in the presence of emodin, the final map is in the EMDB under EMD-22139 and the original micrograph movies and final particle stack is in EMPIAR under EMPIAR-10440. For the high resolution dimeric apo 3a, the final model is in the PDB under 7KJR, the final map is in the EMDB under EMD-22898, and the original micrograph movies and final particle stack is in EMPIAR under EMPIAR-10612.

## Acknowledgements

We thank Dr. Hillel Adesnik for providing emodin and for discussions. We thank Paul Tobias for computational resources at the Cal-Cryo EM facility, and Dr. James Hurley and Dr. Eva Nogales for supporting the microscopy work. We thank Thermo Fisher Scientific for microscope access. We thank Robert Rietmeijer for his gift of TRAAK proteoliposomes. We thank members of the Brohawn lab and Alex Noble for thoughtful feedback on our preprint. Finally, we would like to thank the many people at UC Berkeley and surrounding companies working during the pandemic that helped make this project possible. SGB is a New York Stem Cell Foundation-Robertson Neuroscience Investigator. This work was funded by the New York Stem Cell Foundation, NIGMS grant GM123496, a McKnight Foundation Scholar Award, a Rose Hill Innovator Award, a Sloan Research Fellowship (to SGB), NIGMS grant GM128263 (to DMK), NSF Graduate Research Fellowship DGE1752814 (to SSM), a Howard Hughes Medical Institute Faculty Scholar Award (to DMB), and a Fast Grants Award from Emergent Ventures at the Mercatus Center, George Mason University (to DMB, SGB, and Hillel Adesnik).

## Contributions

BS, DMK, SSM, AK, DMB and SGB conceived of the project. DMK performed all molecular biology, biochemistry, preparation of proteoliposomes, and cryo-EM sample preparation. BS performed all electrophysiology. SSM designed and performed the YO-PRO-1, ruthenium red blocker, and Ca^2+^ flux assays with help from DMB. CMH and DMK processed the cryo-EM data. SS performed light microscopy. JPR and DBT collected cryo-EM data at UC Berkeley and AK collected cryo-EM data at Thermo Fisher Scientific. SGB built and refined the atomic models. DMB and SGB secured funding and supervised research. DMK, BS, CMH, SSM, AK, DMB, and SGB wrote the manuscript with input from all authors.

---

**Figure S1.**
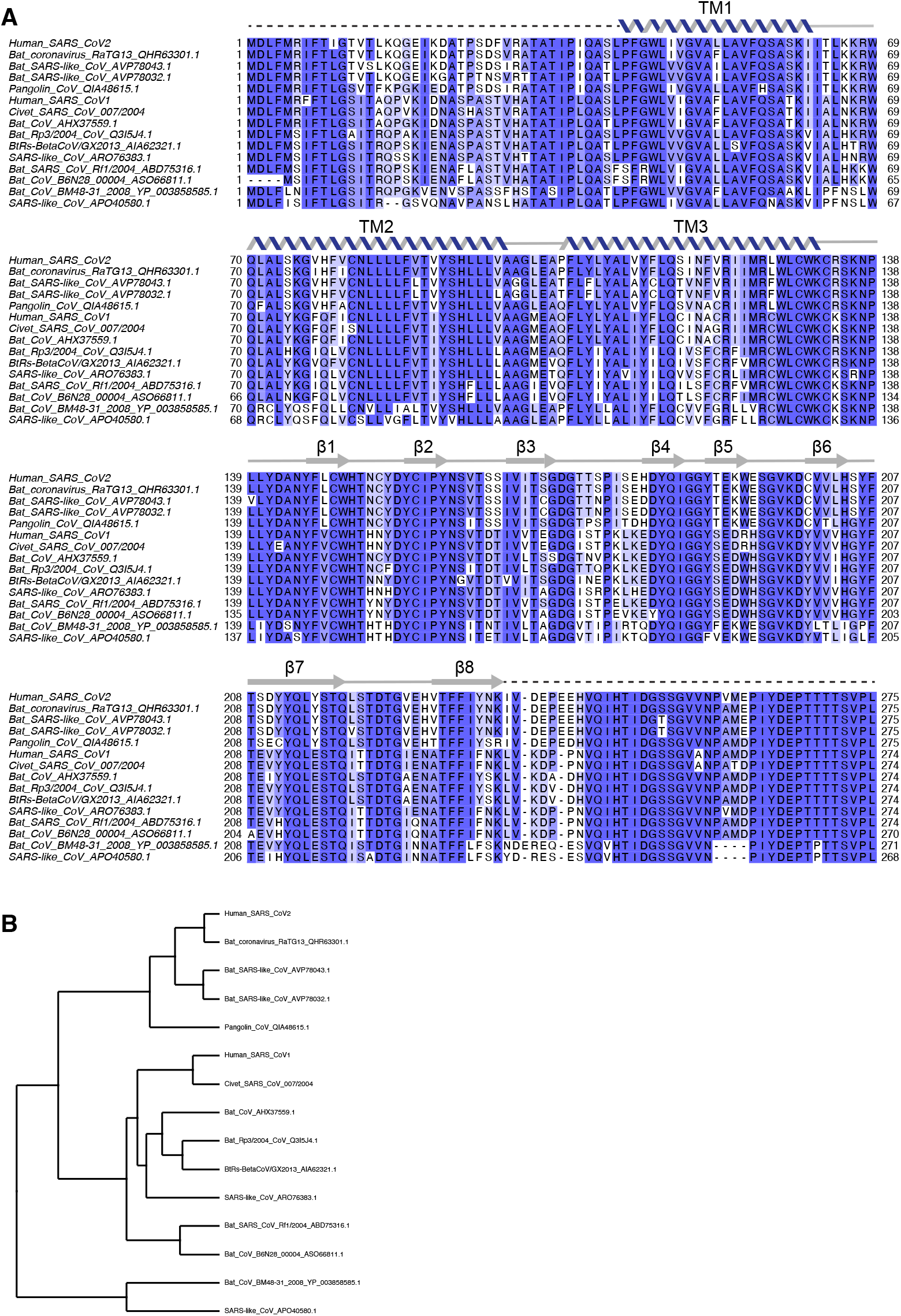
Sequence alignment of 3a from *Betacoronavirus Sarbecovirus*. (A) Alignment of fifteen 3a protein sequences colored by conservation in a ramp from white (not conserved) to dark blue (highly conserved). Accession numbers are indicated. Sequences were selected to maximize diversity among annotated *Sarbecovirus* 3a proteins. Secondary structure for SARS-CoV-2 is drawn above the sequence with unmodeled sequence drawn as dashed lines. (B) Neighbor-joining tree calculated from the alignment in (A).

**Figure S2.**
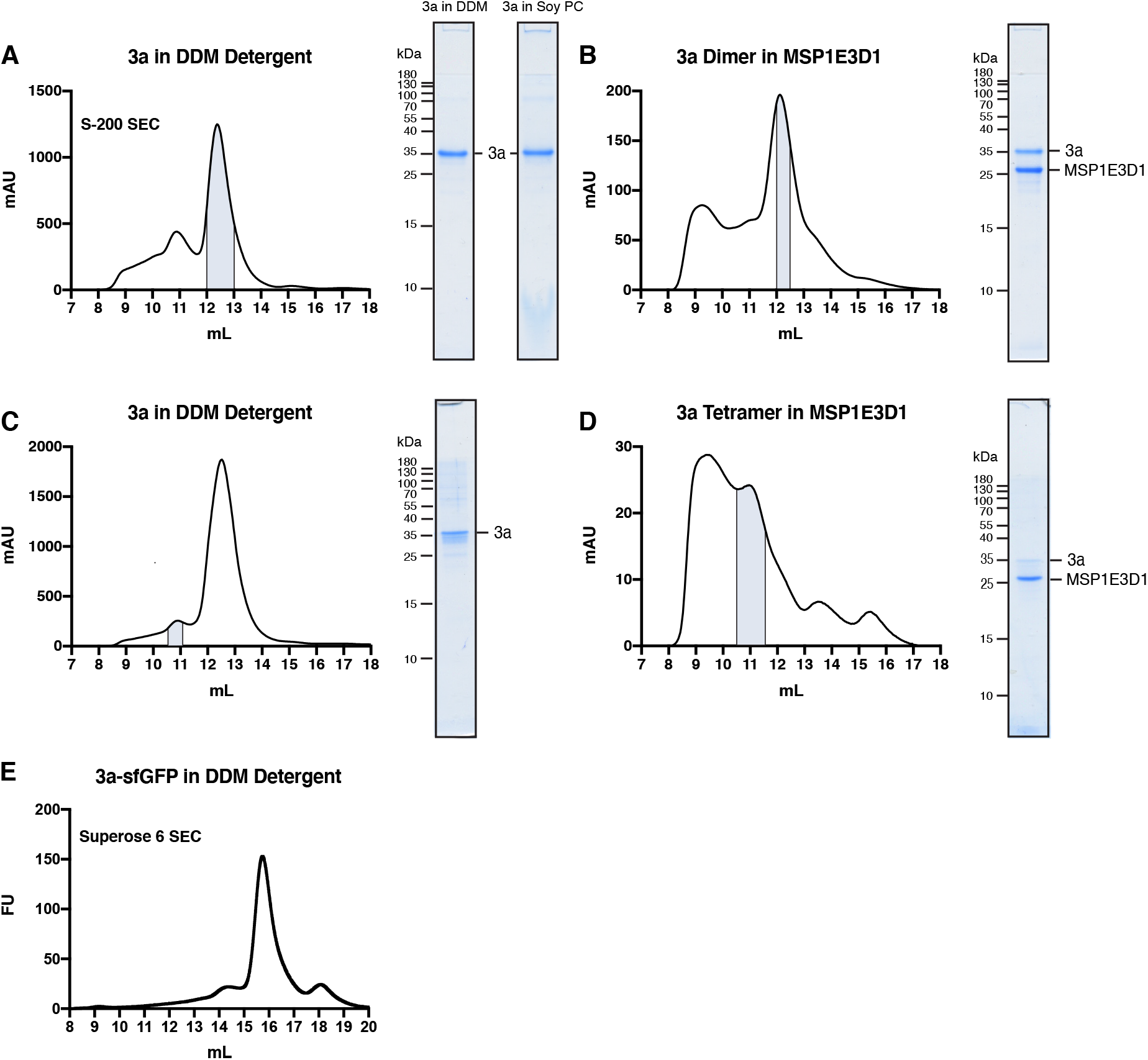
Purification and reconstitution of 3a. (A) Size exclusion chromatogram of 3a expressed in insect cells and extracted and purified in DDM (left). Pooled fractions corresponding to dimeric 3a are highlighted in blue. Coomassie-stained SDS-PAGE of pooled dimeric 3a-containing fractions (center) and of 3a following reconstitution into PC lipids (right). (B) Size exclusion chromatogram of dimeric 3a reconstituted into MSP1E3D1 lipid nanodiscs (left). Pooled fractions are highlighted blue. (C,D) Same as (A,B), but for tetrameric 3a. (E) GFP fluorescence chromatogram of 3a expressed in SF9 cells and extracted in DDM detergent. Samples were run on a Superose 6 column.

**Figure S3.**
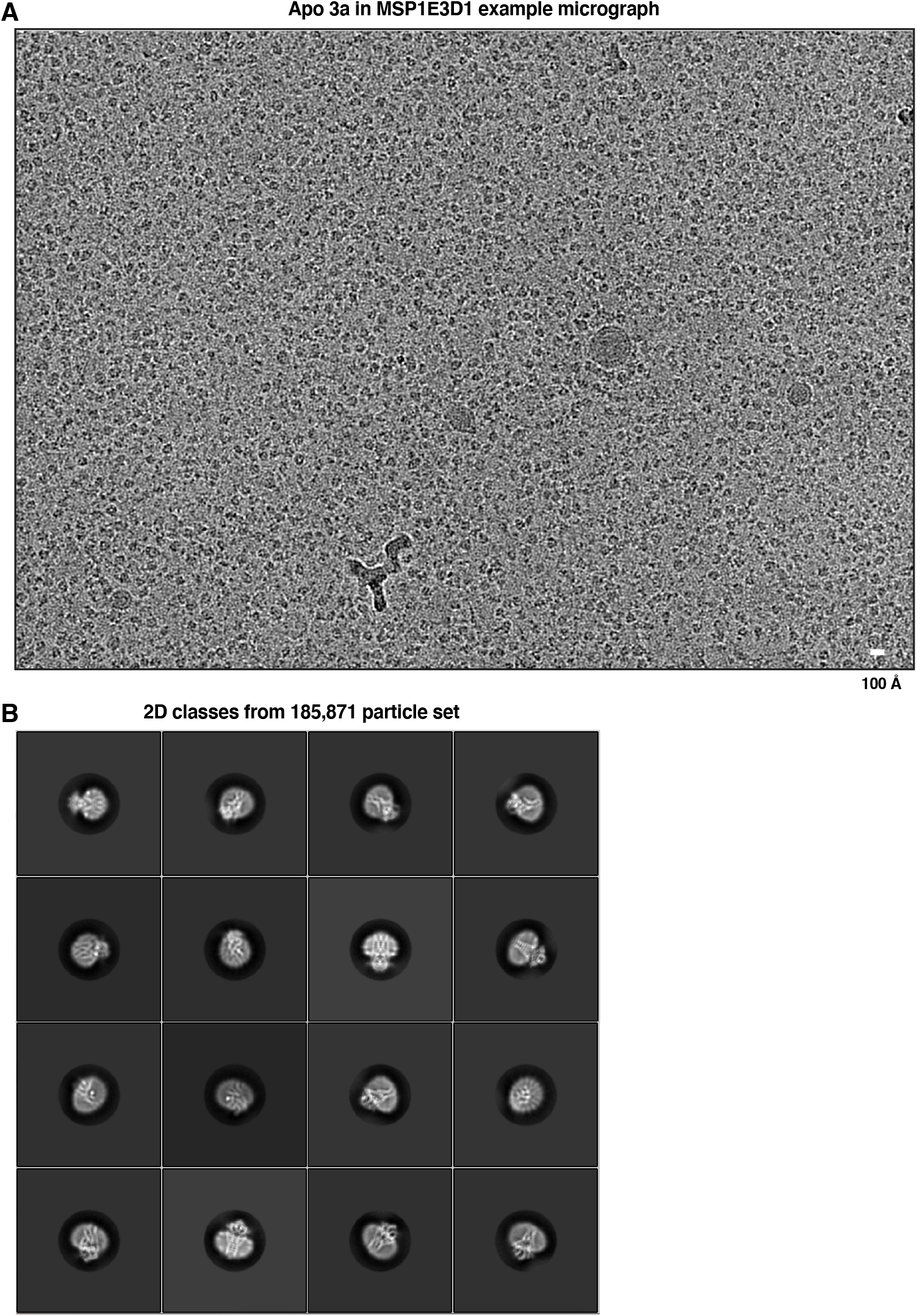
Example micrographs and 2D class averages of dimeric apo 3a in MSP1E3D1 lipid nanodiscs. (A) Representative micrograph and (B) 2D class averages of dimeric apo 3a in MSP1E3D1 lipid nanodiscs.

**Figure S4.**
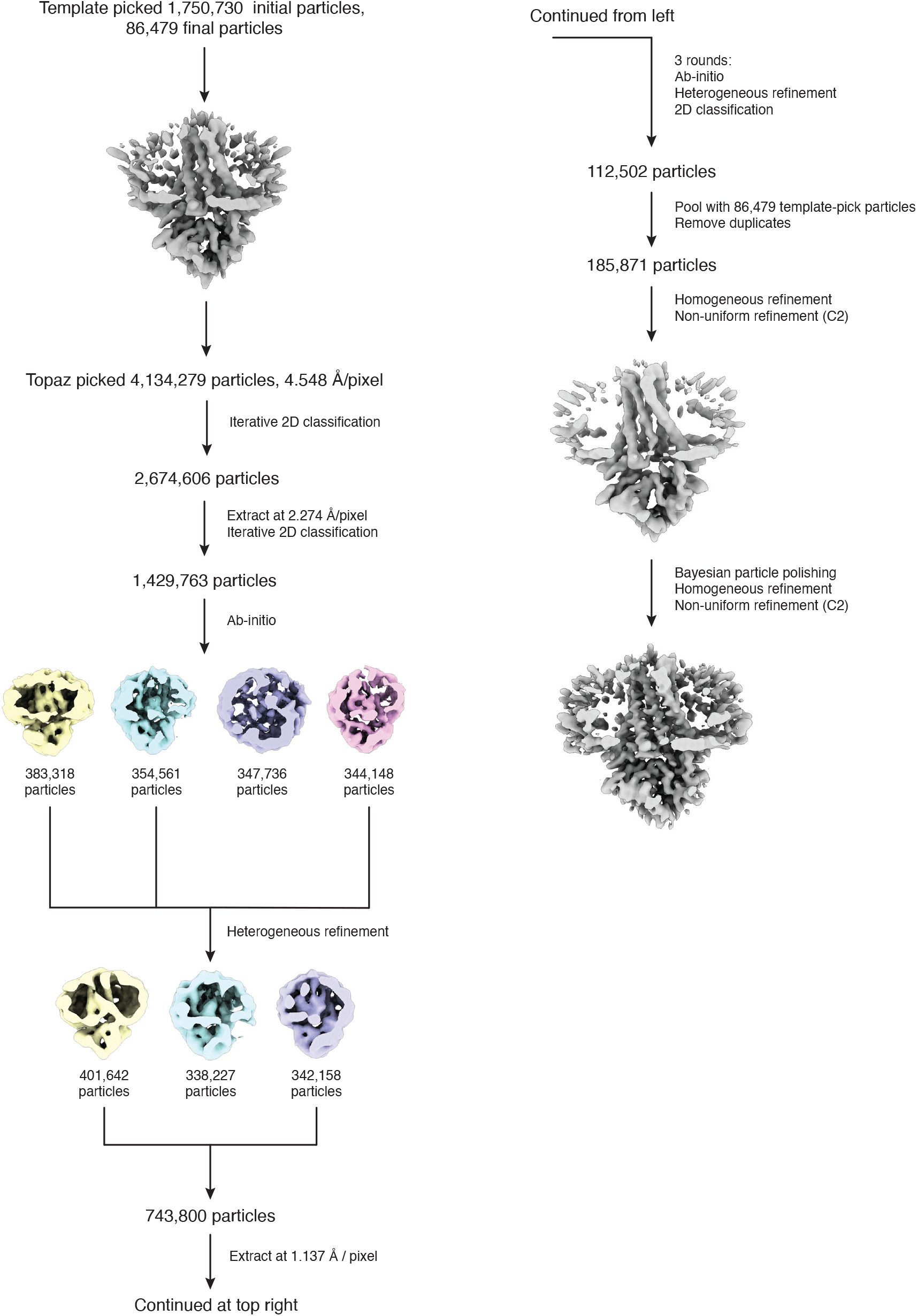
Cryo-EM processing pipeline for dimeric apo 3a in MSP1E3D1 lipid nanodiscs. Overview of Cryo-EM data processing pipeline in cryoSPARC and Relion. See Methods for details.

**Figure S5.**
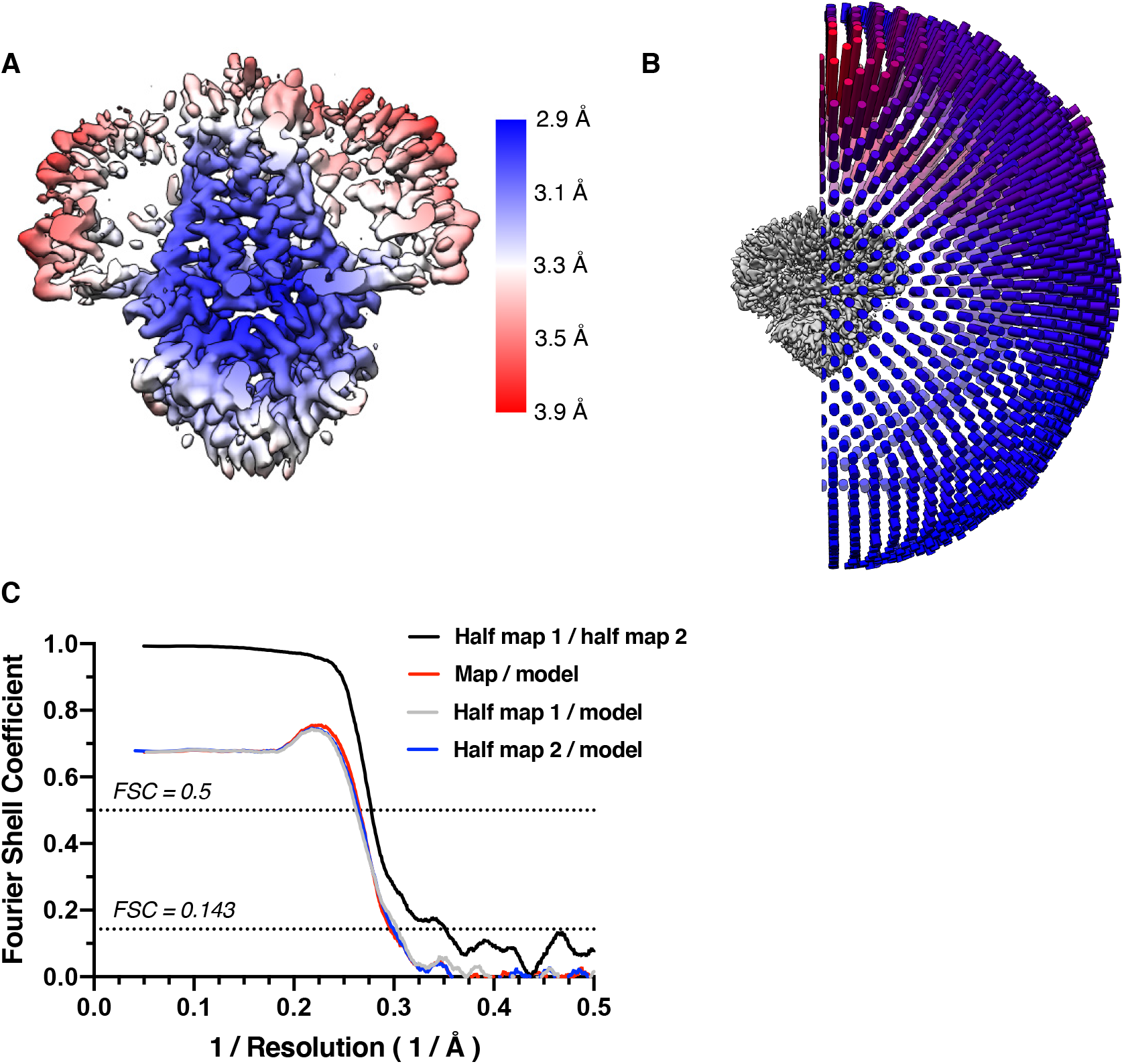
Cryo-EM validation for dimeric apo 3a in MSP1E3D1 lipid nanodiscs. (A) Local resolution estimated in Relion colored as indicated on the final map. (B) Angular distribution of particles used in final refinement with final map for reference. (C) Fourier Shell Correlation (FSC) relationships (unmasked) between (black) the two unfiltered half-maps from refinement and used for calculating overall resolution at 0.143, (red) the final map and model, (gray) half-map one and model, and (blue) half-map and model.

**Figure S6.**
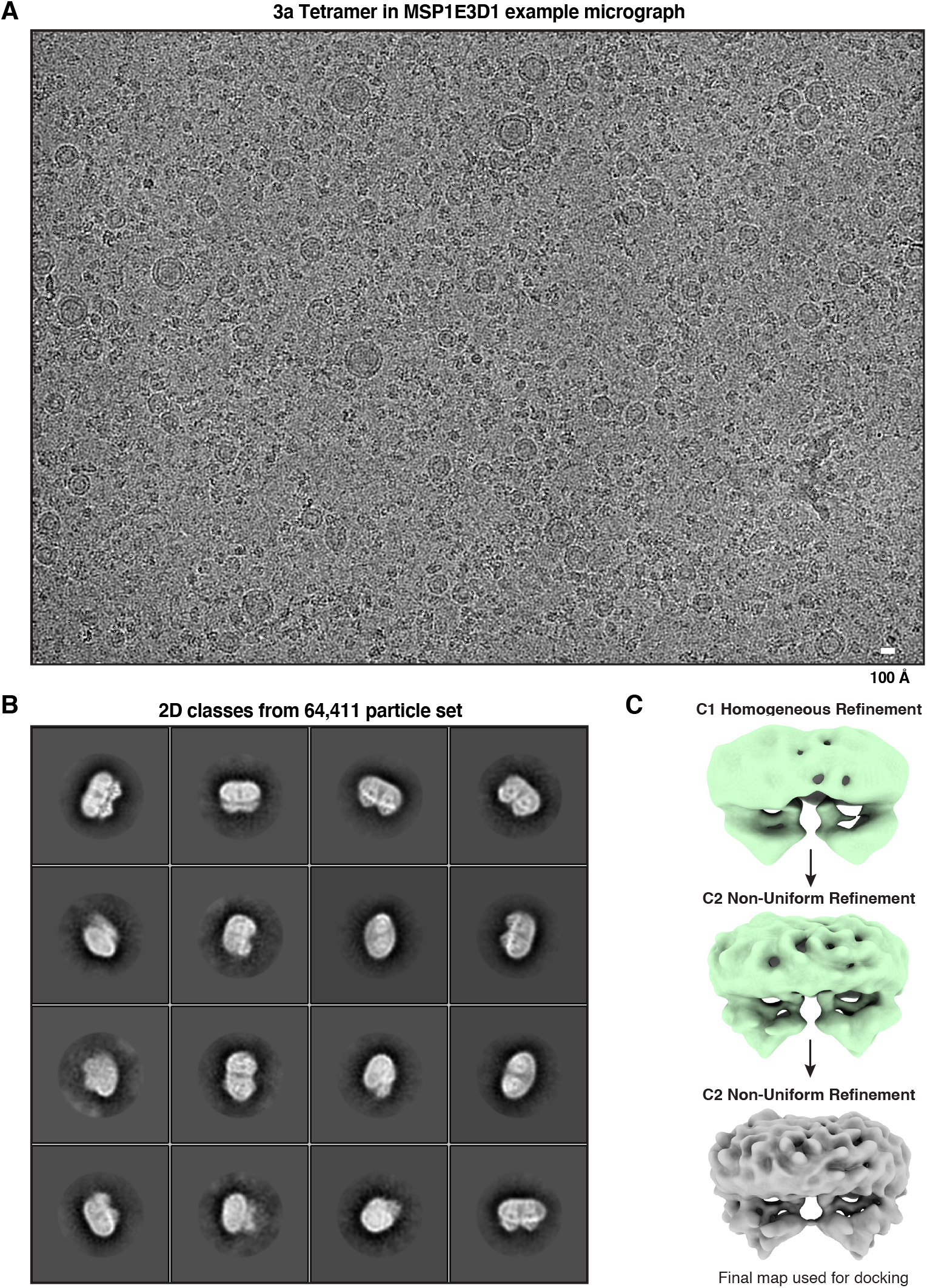
Example micrographs, 2D class averages, and cryo-EM processing pipeline of tetrameric apo 3a in MSP1E3D1 lipid nanodiscs. (A) Representative micrograph and (B) 2D class averages of tetrameric apo 3a in MSP1E3D1 lipid nanodiscs. (C) Map overview pipeline for final steps of processing (Also see Methods).

**Figure S7.**
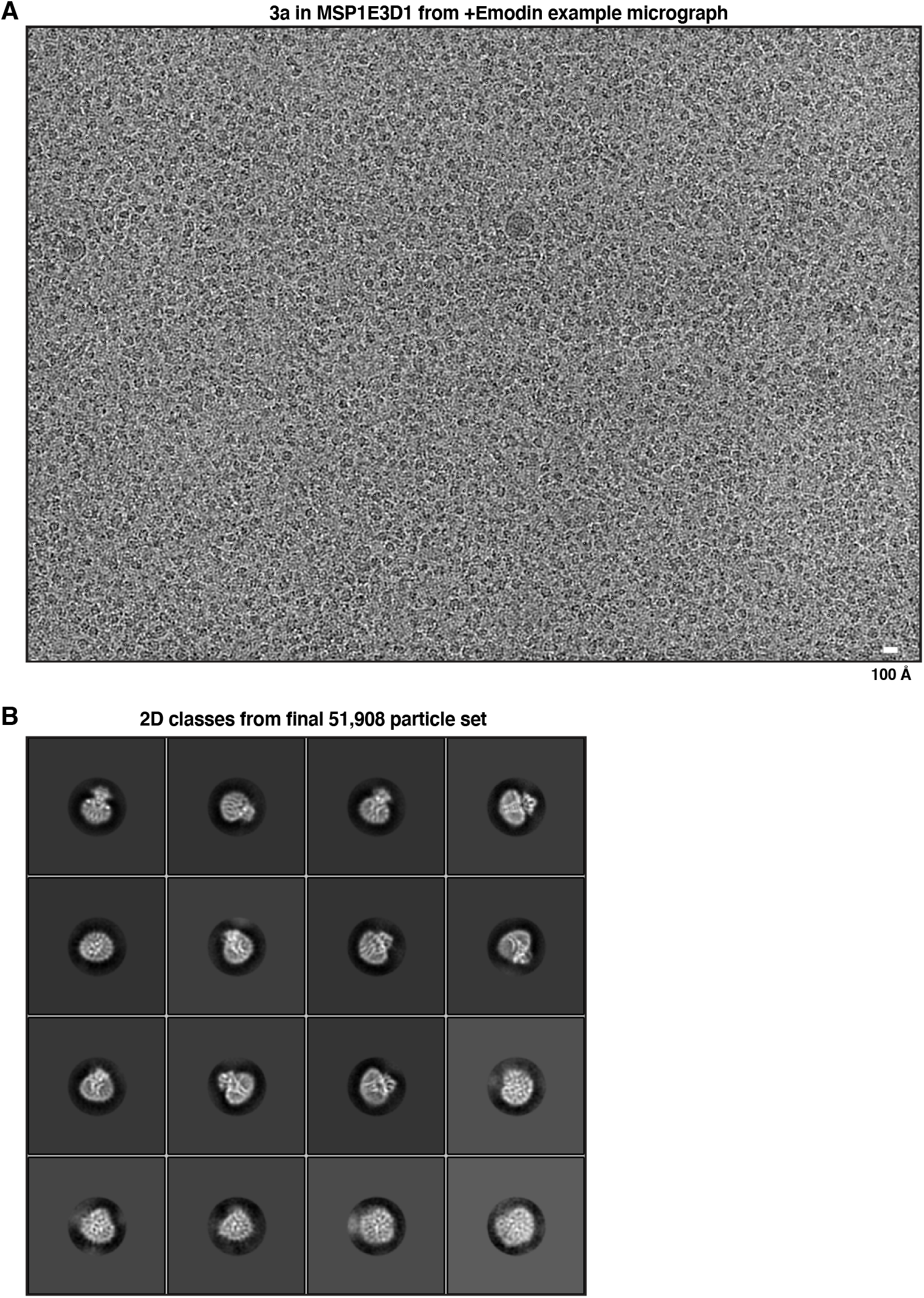
Example micrographs and 2D class averages of dimeric 3a in MSP1E3D1 lipid nanodiscs with emodin added. (A) Representative micrograph and (B) 2D class averages of dimeric 3a in MSP1E3D1 lipid nanodiscs with emodin added.

**Figure S8.**
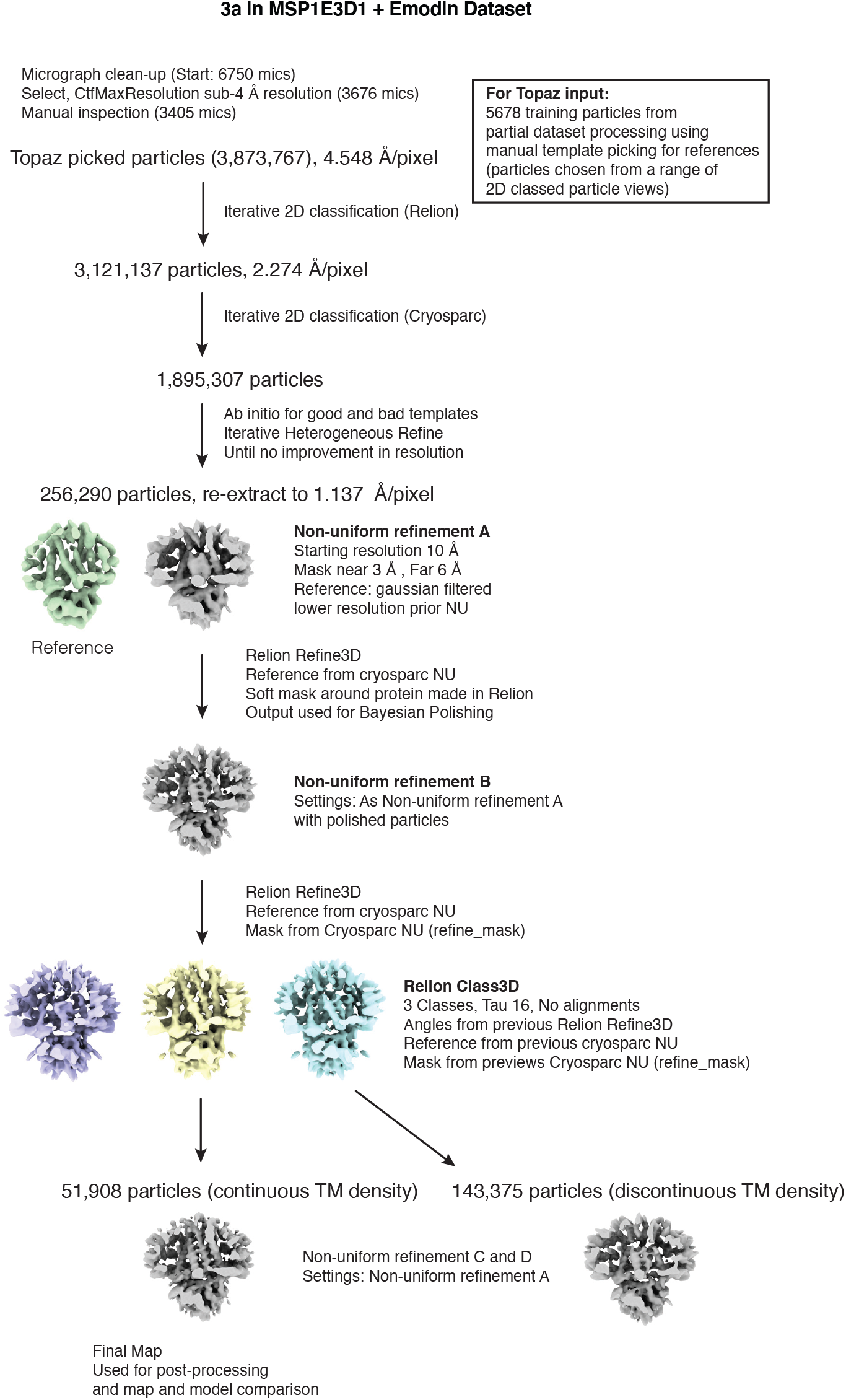
Cryo-EM processing pipeline for dimeric 3a in MSP1E3D1 lipid nanodiscs with emodin added. Overview of Cryo-EM data processing pipeline in cryoSPARC and Relion. See Methods for details.

**Figure S9.**
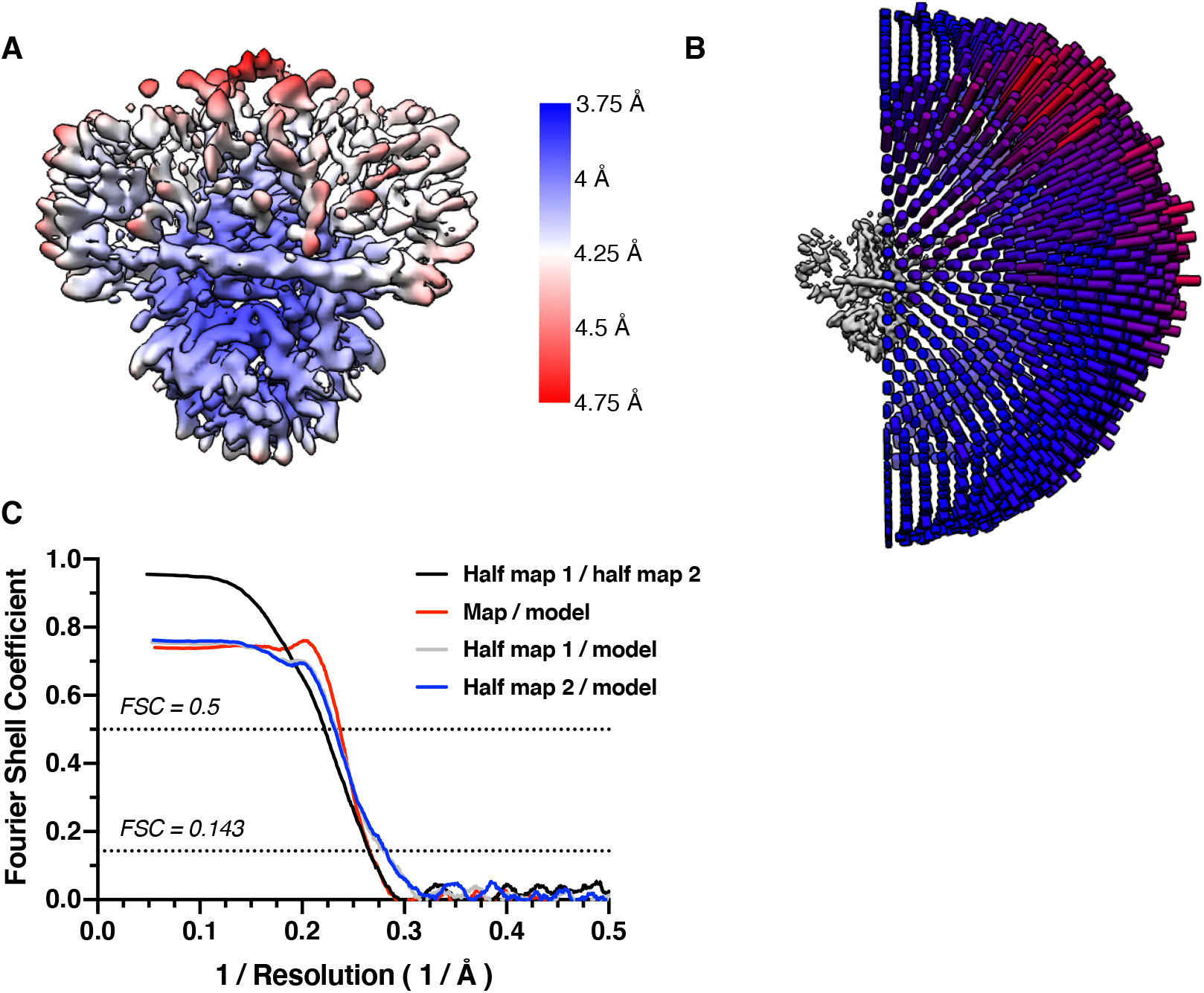
Cryo-EM validation for dimeric 3a in MSP1E3D1 lipid nanodiscs with emodin added. (A) Local resolution estimated in Relion colored as indicated on the final map. (B) Angular distribution of particles used in final refinement with final map for reference. (C) Fourier Shell Correlation (FSC) relationships (masked) between (black) the two unfiltered half-maps from refinement and used for calculating overall resolution at 0.143, (red) the final map and model, (gray) half-map one and model, and (blue) half-map and model.

**Figure S10.**
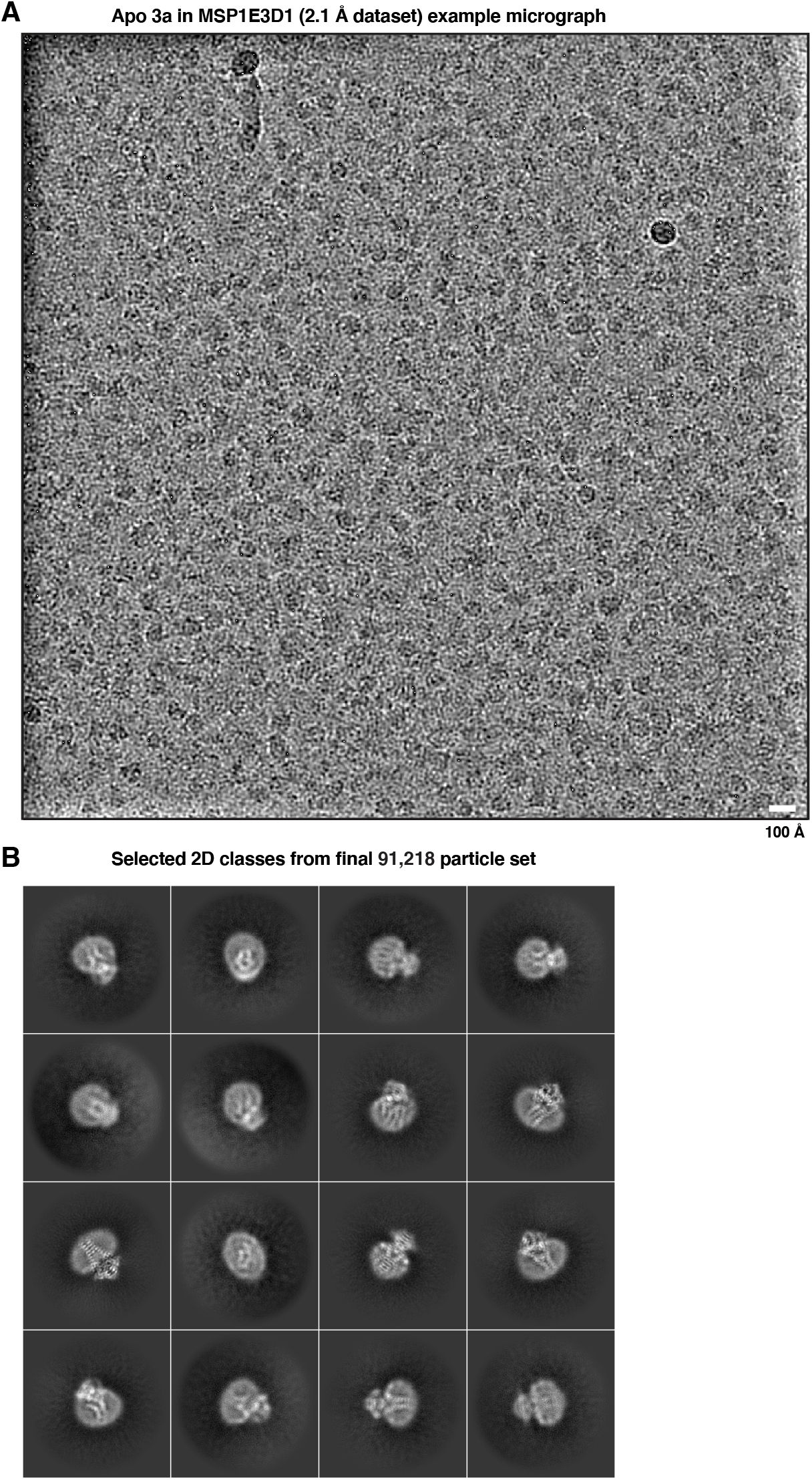
Example micrographs and 2D class averages of dimeric apo 3a in MSP1E3D1 lipid nanodisc collected on the Krios with CFEG, Selectris, and Falcon 4. (A) Representative micrograph and (B) 2D class averages of dimeric apo 3a in MSP1E3D1 lipid nanodiscs from cryoSPARC.

**Figure S11.**
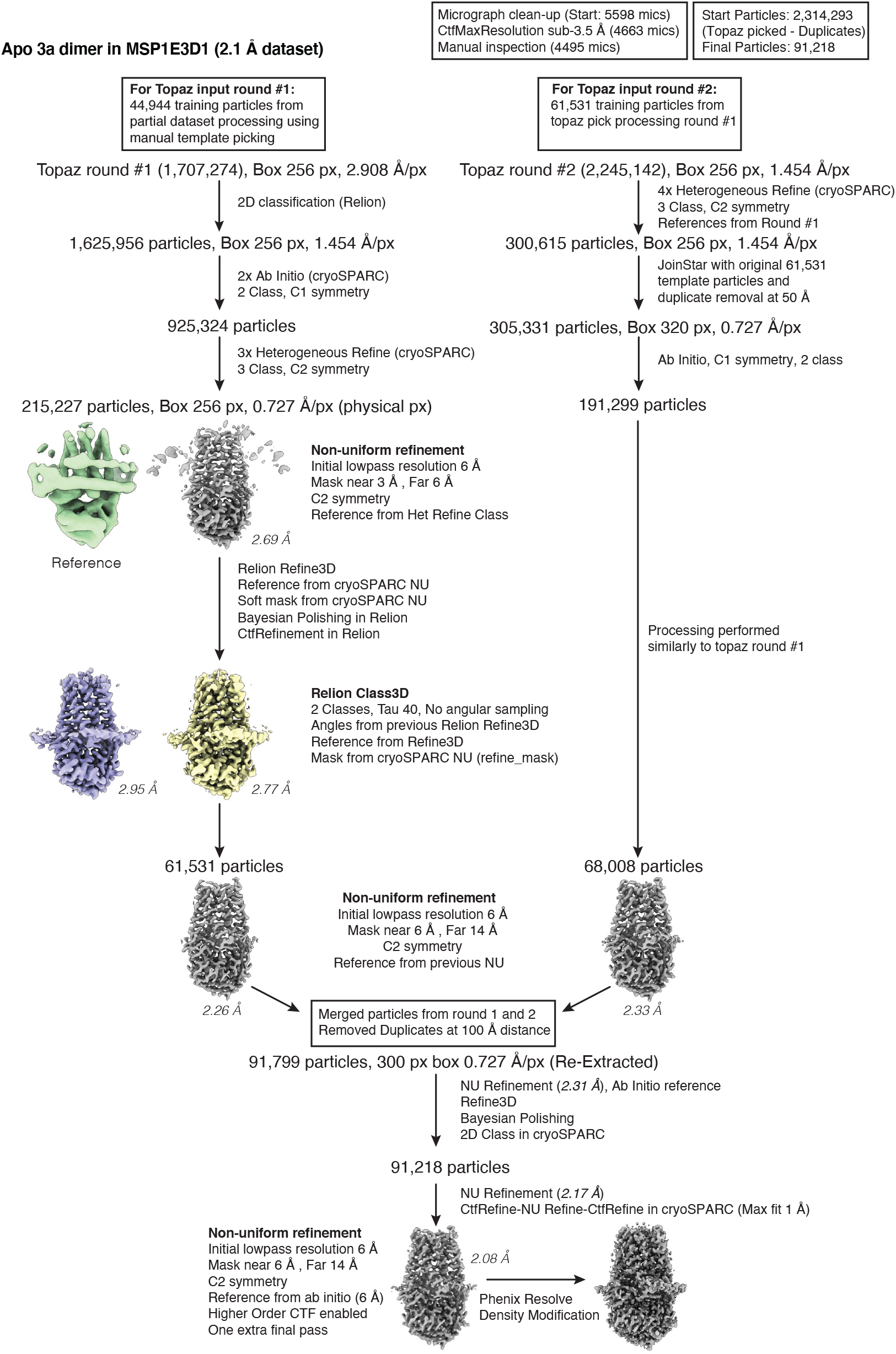
Cryo-EM processing pipeline for dimeric apo 3a in MSP1E3D1 lipid nanodiscs. Overview of Cryo-EM data processing pipeline in cryoSPARC and Relion. See Methods for details.

**Figure S12.**
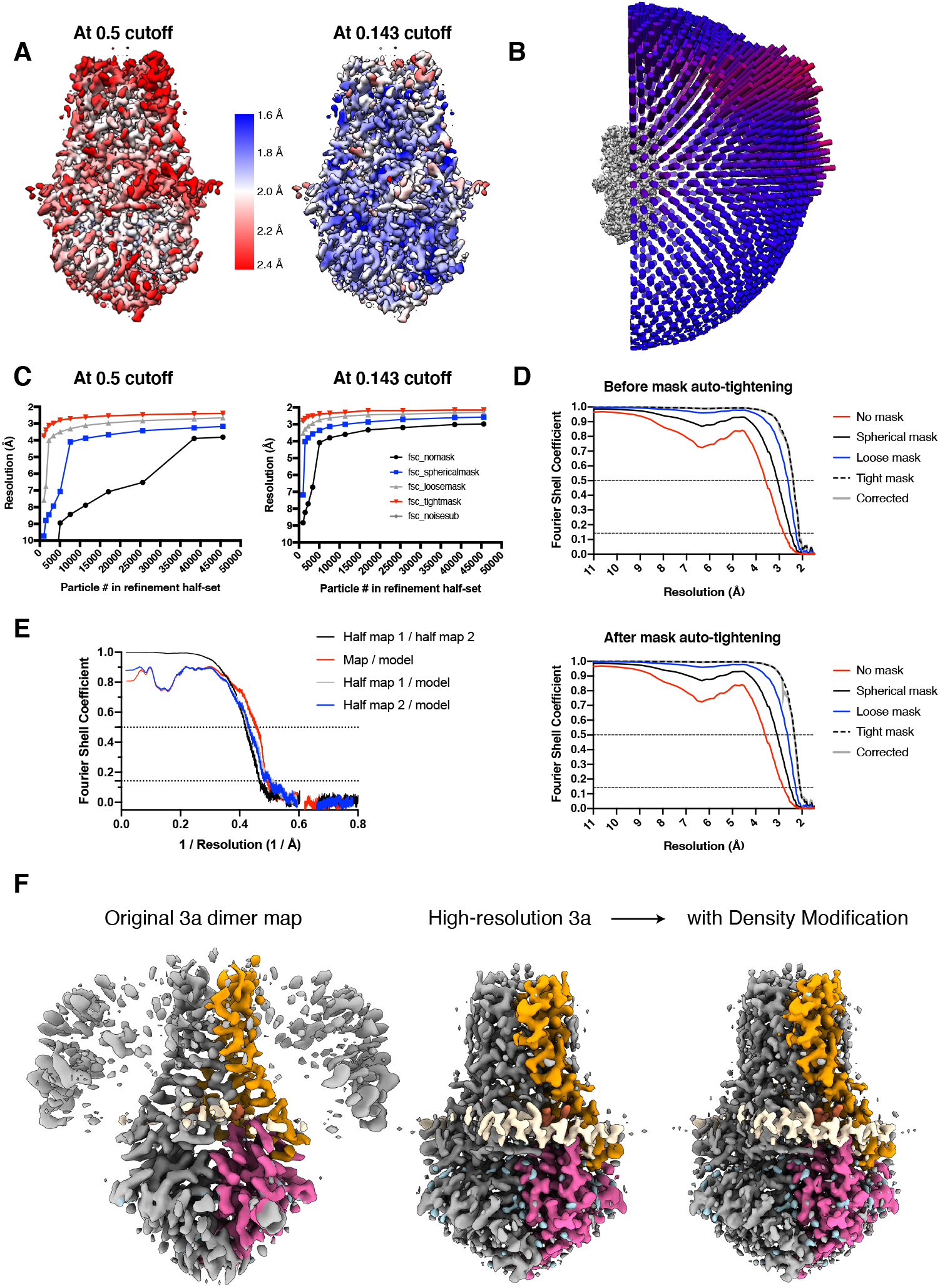
Cryo-EM validation for dimeric apo 3a in MSP1E3D1 lipid nanodiscs. (A) Local resolution estimated in cryoSPARC at the indicated FSC thresholds colored as indicated on the density modified map. (B) Angular distribution of particles used in final refinement with map for reference. (C) ResLog analysis conducted in cryoSPARC at the indicated FSC thresholds. (D) Fourier Shell Correlation as calculated in cryoSPARC before (top) and after (bottom) mask auto-tightening in the final round of refinement. (E) Fourier Shell Correlation (FSC) relationships (masked) calculated in Phenix between (black) the two unfiltered half-maps from refinement and used for calculating overall resolution at 0.143, (red) the final map and model, (gray) half-map one and model, and (blue) half-map and model. (F) Comparison of the original (2.9 Å) 3a map left to the high-resolution (2.1 Å) map before (middle) and after (right) Phenix density modification.

**Figure S13.**
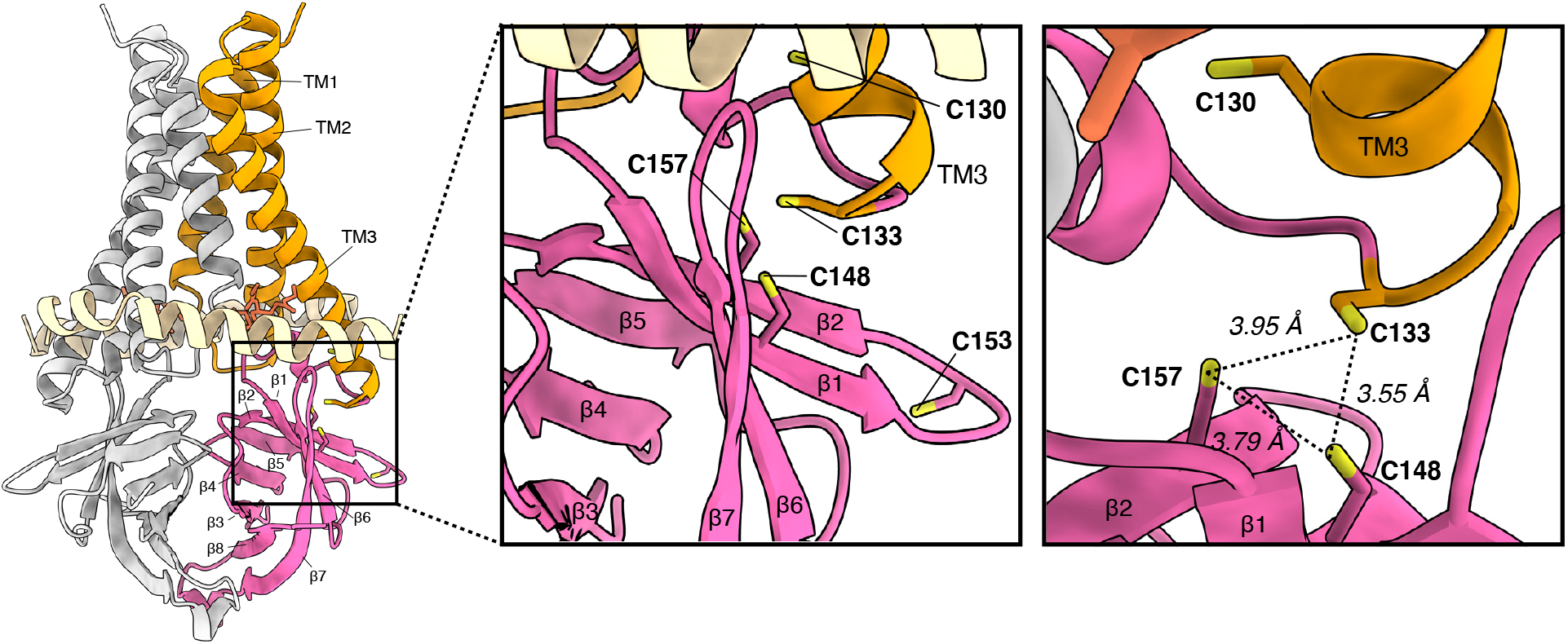
A cysteine rich pocket in 3a. Full model (left) with boxed region zoom-in (middle) and an alternate view (right) to show the cysteine-rich region of 3a. Distances (dotted lines) between the reduced cysteines are displayed.

**Figure S14.**
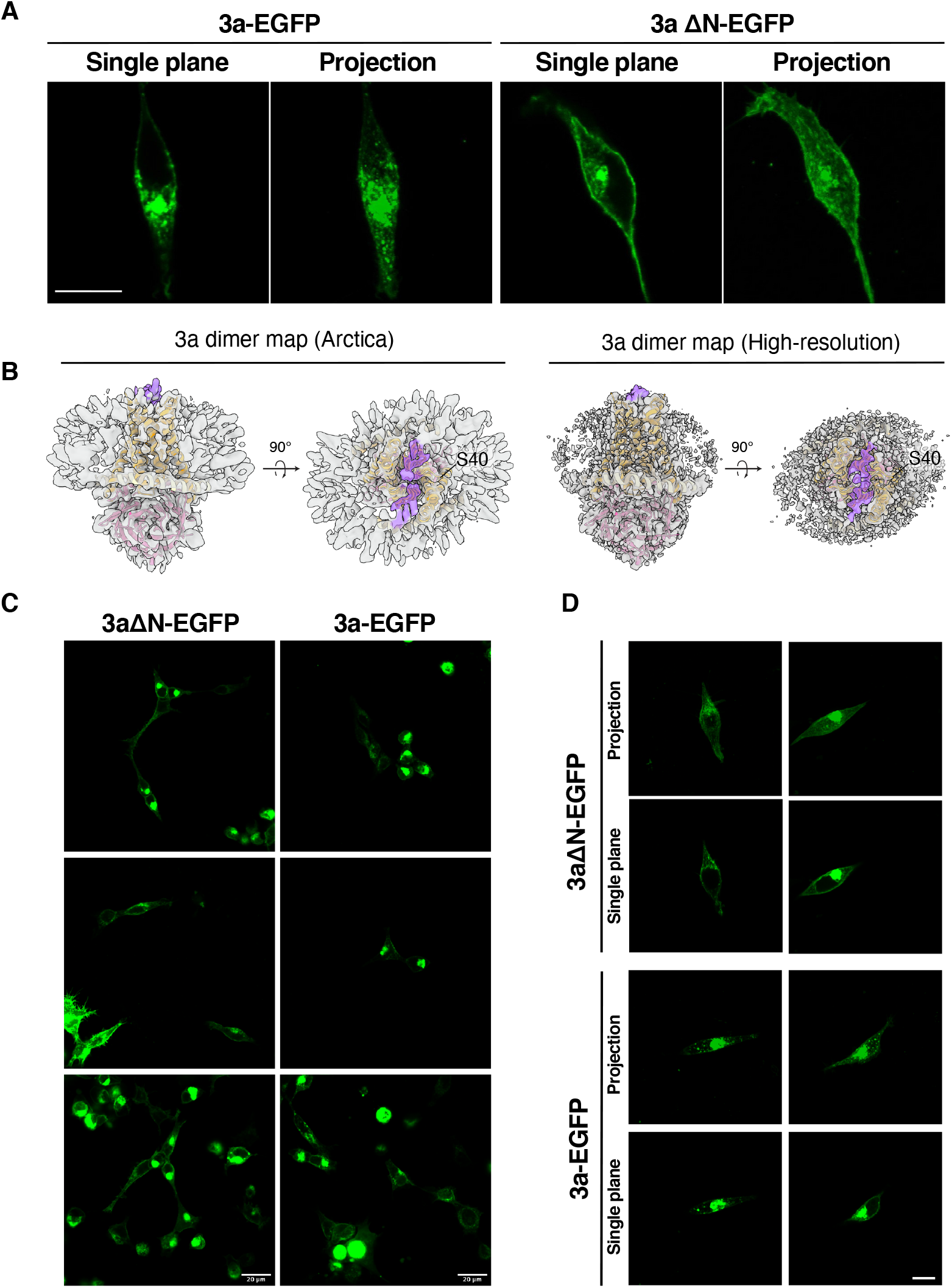
Localization of 3a-EGFP and 3aΔN-EGFP expressed in HEK cells. (A) 3a-GFP fluorescence localization in HEK293 cells transfected with 3a-EGFP or 3aΔN-EGFP. Single plane and brightest-point projection are displayed for each. Scale bar, 10 μm. (B) Side view and view from the extracellular/lumenal space for dimeric 3a cryo-EM density (gray) from the original (left) and high-resolution (right) maps with unmodeled extended density above the mouth of the pore that may correspond to the N-terminal regions colored in purple. A 3a dimer model is drawn orange (TMD) and pink (CD) inside the density. The position of the final modeled N-terminal residue (S40) is indicated. (C) 3a-EGFP or 3aΔN-EGFP field of view with multiple cells imaged using a 20X objective. Scale bar, 20 μm. (D) Additional images of cells imaged with the 63X objective with both single plane and brightest-point projections displayed. Scale bar, 10 μm.

**Figure S15.**
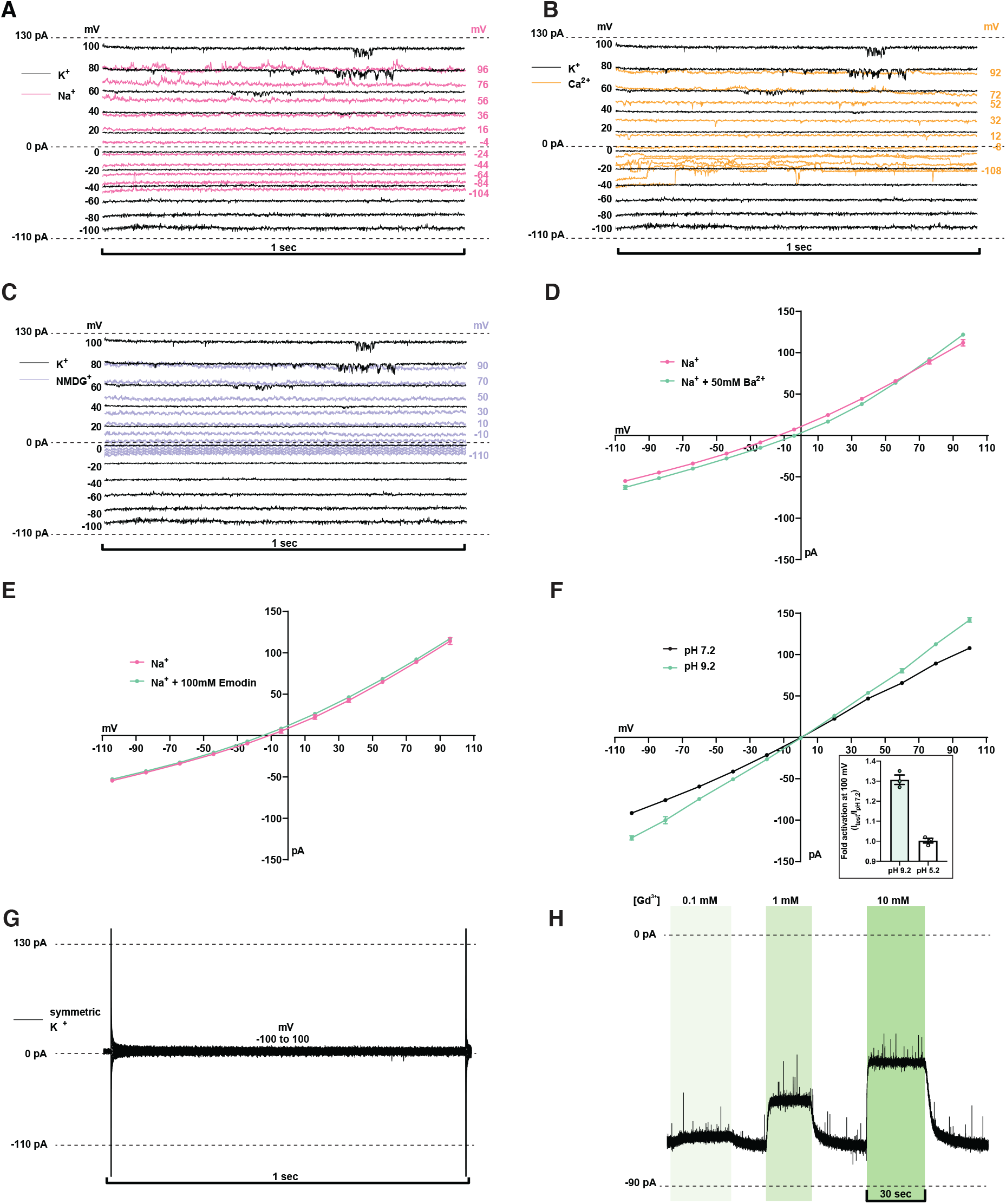
Patch recordings from 3a-proteoliposomes. (A-C,G) Representative current recordings from 3a-proteoliposome. Currents were recorded with the following protocol: V_hold_ = 0 mV, V_test_ = −100 to 100 mV, ΔV = 20 mV, t_test_=1 sec. Voltages indicated were corrected after recording for liquid junction potential. (A) 150 mM K^+^ (black) or 150 mM Na^+^ (pink) bath solution. (B) 150 mM K^+^ (black) or 75 mM Ca^2+^ (orange) bath solution. (C) K^+^ (black) or 150 mM NMDG^+^ (blue) bath solution. (D) Ba^2+^ does not block 3a currents. Current-voltage relationship plotted from a recording in 150 mM Na^+^ (pink) or 150 mM Na^+^ with 50 mM Ba^2+^ (green) bath solution. (E) Emodin does not block 3a currents. Current-voltage relationship plotted from a recording in 150 mM Na^+^ (pink) or 150 mM Na^+^ with 100 μM emodin (green) bath solution. (F) pH sensitivity of 3a. Current-voltage relationship plotted from a recording in 150 mM K^+^ pH 7.2 (black) and 150 mM K^+^ pH 9.2 (green) bath solution. (inset) Fold activation at +100 mV at pH 9.2 or pH 5.2 compared to pH 7.2. (G) Representative current recordings from mock reconstituted (empty) liposome patch. (H) Gap-free current recording in symmetric 150 mM KCl held at −80 mV during bath solution exchange and washout of varying Gd^3+^ concentrations represented by vertical green bars.

**Figure S16.**
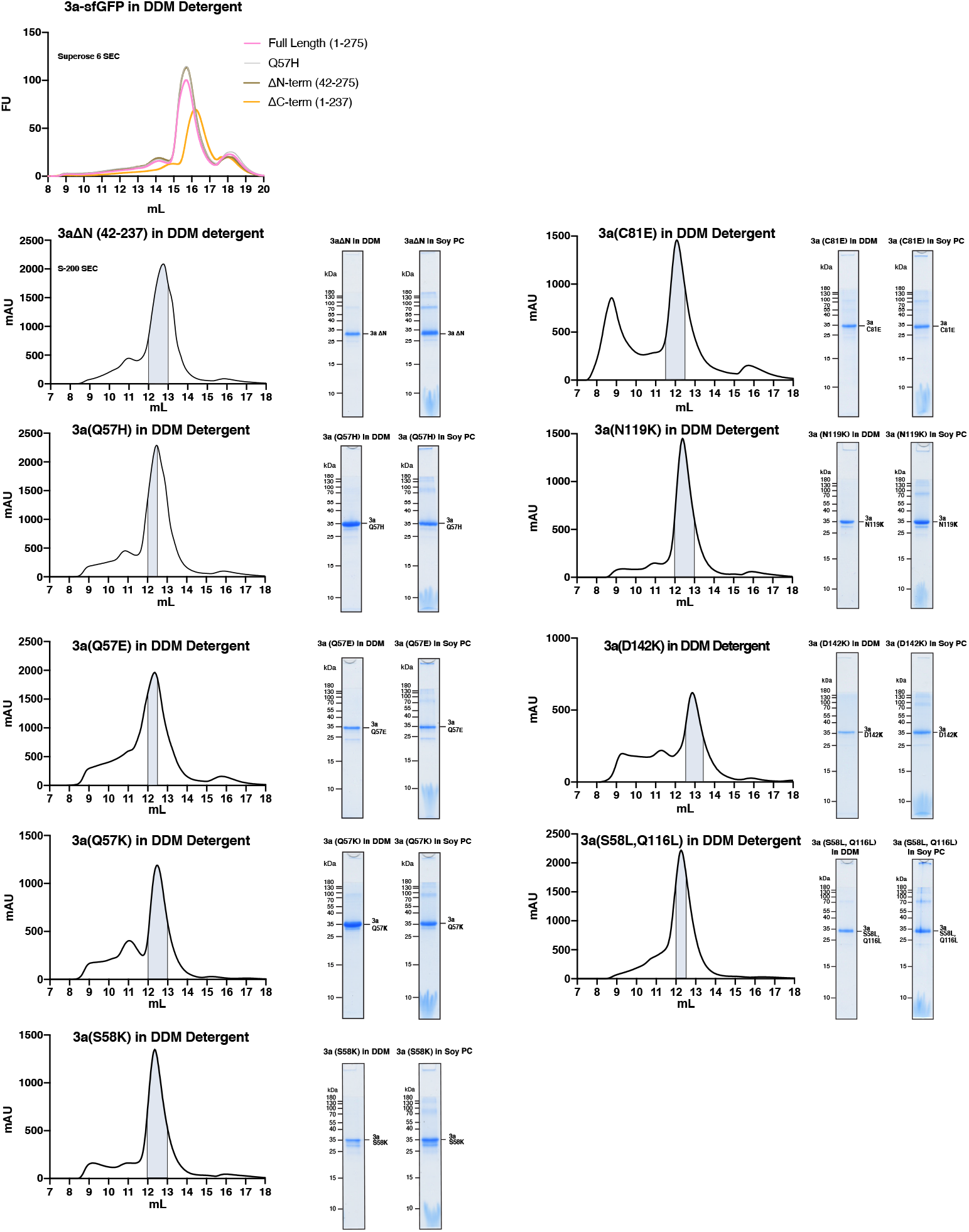
Purification and liposome reconstitution of 3a mutants and truncation. Top: GFP fluorescence chromatogram (FSEC) of 3a, as well as indicated mutants and truncations, expressed in SF9 cells and extracted in DDM detergent. Samples were run on a Superose 6 column. For all other panels: Size exclusion chromatogram from a s200 column of indicated 3a constructs expressed in insect cells and extracted and purified in DDM (left), coomassie-stained SDS-PAGE of pooled dimeric 3a construct -containing fractions (center), and of 3a following reconstitution into PC lipids (right).

**Figure S17.**
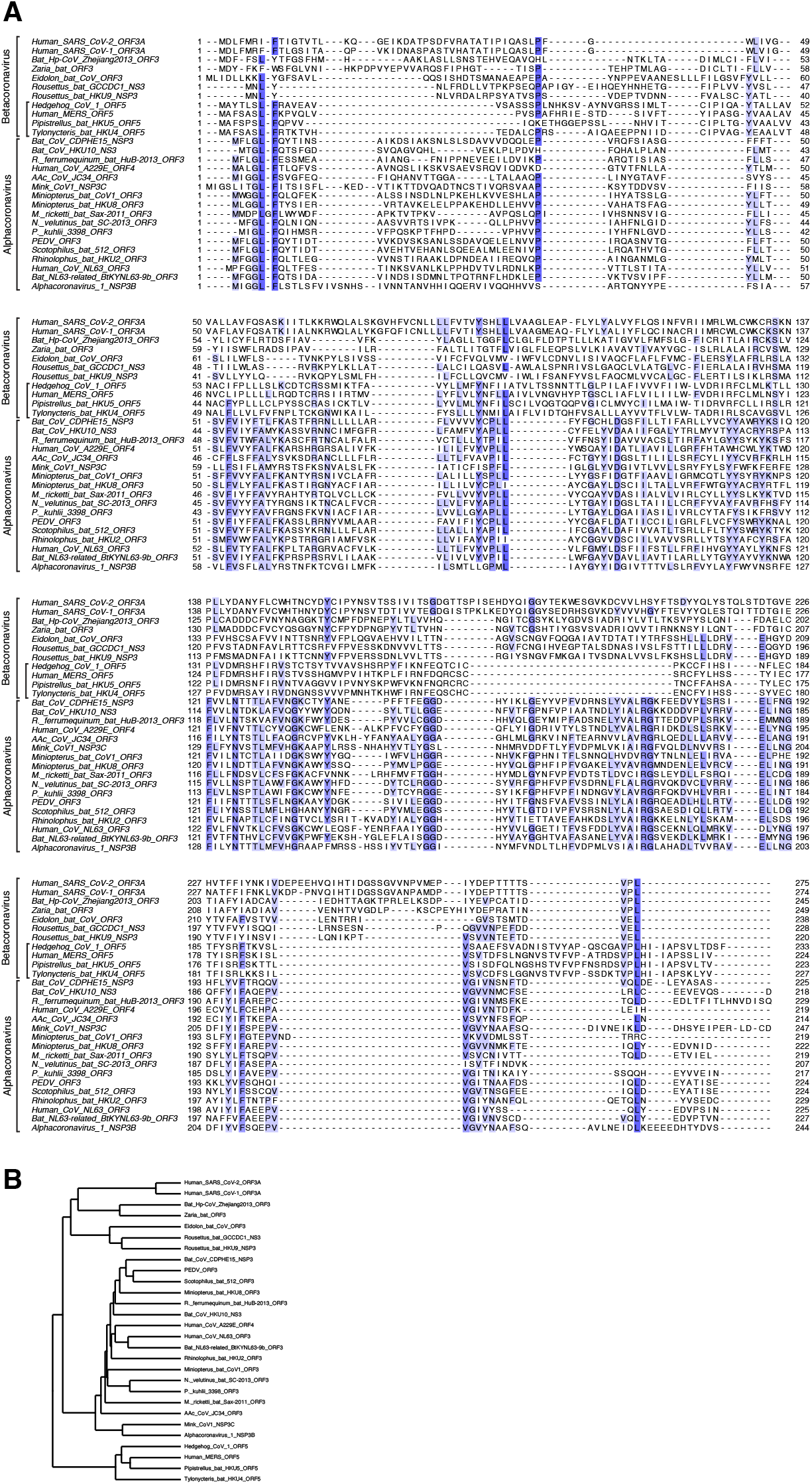
Sequence alignment of 3a-like proteins across *Coronaviridae*. (A) Alignment of twenty-eight 3a-like protein sequences colored by conservation in a ramp from white (not conserved) to dark blue (highly conserved). Accession numbers are listed in Table 2. Sequences were selected from representative species from each Coronavirus subgenus. *Alphacoronavirus* and Betacoronavirus clades are indicated. Within Betacoronavirus the subgenus *Merbecovirus* is also indicated with a bar. (B) Neighbor-joining tree calculated from the alignment in (A).

**Table S1.** Cryo-EM data collection, processing, refinement, and modeling data.

| Data collection | Dimeric apo 3a<br>(2.9 Å) | Dimeric 3a<br>with emodin | Tetrameric<br>apo 3a | Dimeric<br>apo 3a<br>(2.1 Å) |
| --- | --- | --- | --- | --- |
| PDB | 6XDC | n/a | n/a | 7KJR |
| EMDB | 22136 | 22139 | 22138 | 22898 |
| EMPIAR | 10439 | 10440 | 10441 | 10612 |
| Total movies # | 6309 | 6750 | 7092 | 5598 |
| Selected movies # | 2595 | 3405 | 4324 | 4495 |
| Magnification | 36,000 x | 36,000 x | 36,000 x | 165,000 x |
| Voltage (KV) | 200 | 200 | 200 | 300 |
| Electron exposure (e<sup>-</sup>/Å<sup>2</sup>) | 50.325 or 53.72<br>(1-2007) (2008-6309) | 47.21 | 49.95 | 50 |
| Frame # | 50 | 50 | 50 | 1449 or<br>1379 (EER) |
| Defocus range (um) | -0.6 to -2.0 | -0.6 to -2.0 | -0.6 to -2.0 | -0.5 to -1.1 |
| Super resolution pixel size (Å<sup>2</sup>) | 0.5685 | 0.5685 | 0.5685 | 0.3685 |
| Binned pixel size (Å<sup>2</sup>) | 1.137 | 1.137 | 1.137 | 0.727 |
| **Processing** |  |  |  |  |
| Initial particle images (no.) | 4,134,279 | 3,873,767 | 1,282,913 | 2,314,293 |
| Final particle images (no.) | 185,871 | 51,908 | 64,410 | 91,218 |
| Map resolution Masked (Å, FSC = 0.143) | 2.9 | 3.69 | 6.5 | 2.08 |
| Symmetry imposed | C2 | C2 | C2 | C2 |
| **Refinement** |  |  |  |  |
| Model resolution (Å, FSC = 0.143 / FSC = 0.5) | 3.2/3.6 |  |  | 2.0 / 2.2 |
| Map-sharpening B factor (Å<sup>2</sup>) | -111.2 |  |  | -43.5 |
| Composition |  |  |  |  |
| Number of atoms | 3150 |  |  | 3840 |
| Number of protein residues | 386 |  |  | 448 |
| Number of ligands |  |  |  | 2 |
| Number of waters |  |  |  | 122 |
| R.m.s. deviations |  |  |  |  |
| Bond lengths (Å) | 0.006 |  |  | 0.005 |
| Bond angles (Å) | 0.785 |  |  | 0.729 |
| Validation |  |  |  |  |
| MolProbity score | 1.55 |  |  | 1.42 |
| Clashscore | 4.63 |  |  | 7.60 |
| Ramachandran plot |  |  |  |  |
| Favored (%) | 96.03 |  |  | 98.17 |
| Allowed (%) | 3.97 |  |  | 1.13 |
| Disallowed (%) | 0 |  |  | 0 |
| Rotamer outliers (%) | 1.15 |  |  | 0.25 |
| Mean B factor (Å<sup>2</sup>) |  |  |  |  |
| Protein | 108.61 |  |  | 23.30 |
| Ligand |  |  |  | 51.72 |
| Water |  |  |  | 22.11 |

**Table S2.**
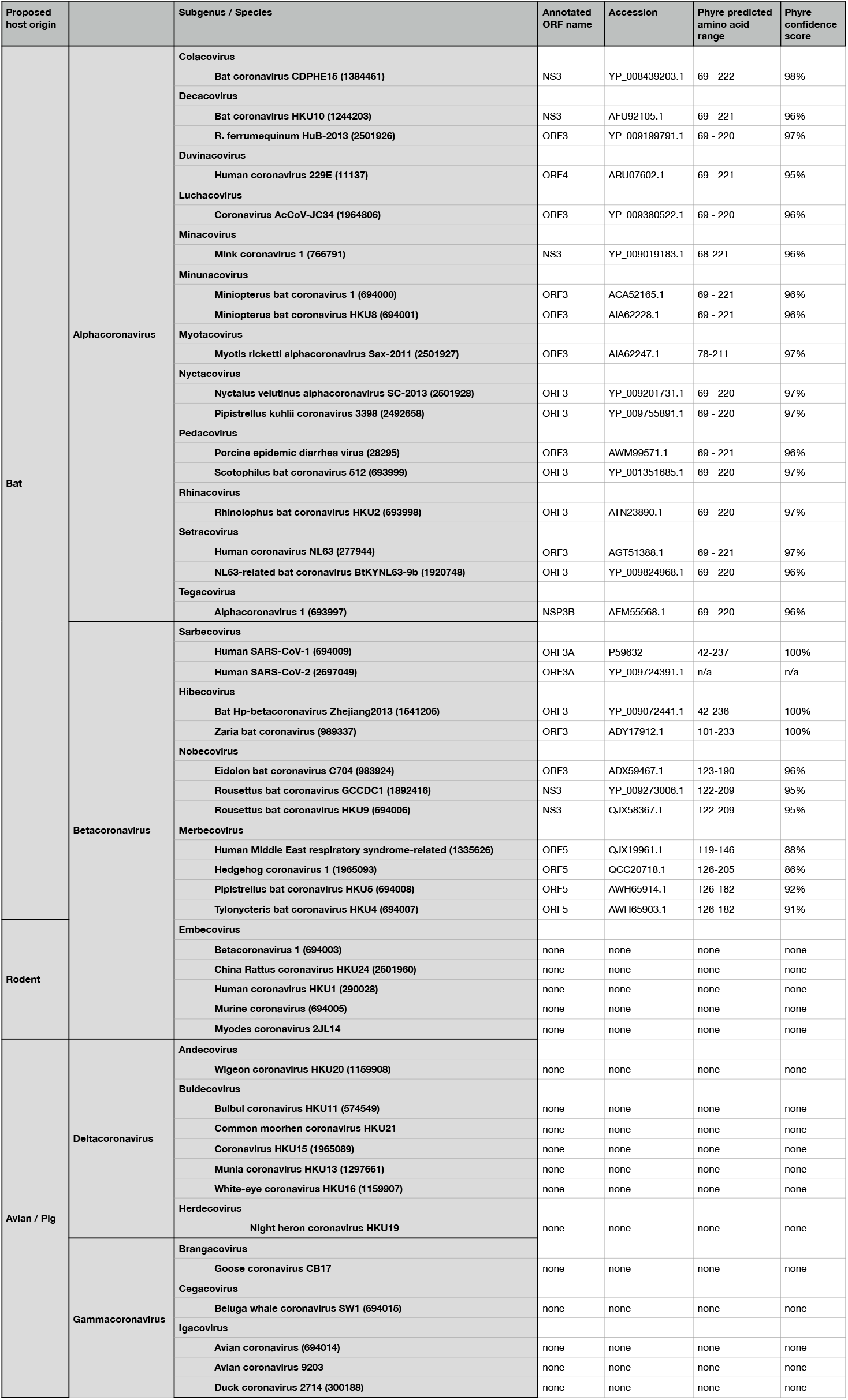
3a homologs across *Coronaviridae*.

